# Modeling Immunosenescence on-a-chip: a platform for cancer vaccine efficacy assessment

**DOI:** 10.1101/2025.05.14.654059

**Authors:** Surjendu Maity, Alireza Hassani Najafabadi, Satoru Kawakita, Danial Khorsandi, Can Yilgor, Christopher Jewell, Neda Mohaghegh, Mehmet Remzi Dokmeci, Ali Khademhosseini, Vadim Jucaud

## Abstract

Immunosenescence dramatically reduces cancer vaccine efficacy in elderly patients, who represent the majority of cancer cases. Despite this clinical reality, age-related immune decline is rarely considered in preclinical testing. Therefore, novel in vitro models to test cancer vaccine efficacy, considering immunosenescence, are needed. Our novel lymph node paracortex on-a-chip (LNPoC) platform addresses this gap by recapitulating age-dependent immune responses against cancer vaccines, specifically antigen presentation, antigen-specific T cell activation, and antitumoral responses. Using this platform, we demonstrated that bone marrow-derived antigen-presenting cells (APCs) from young mice (6-7 weeks) displayed significantly enhanced ovalbumin (OVA) peptide presentation compared to APCs from older mice (35-36 weeks). This age-dependent difference translated to significantly greater OVA-specific CD8+ T cell activation and increased cytotoxicity against B16-OVA cancer cells. These age-dependent differences are unique to our LNPoC and undetectable in traditional 2D cultures, confirming that our LNPoC was more effective than 2D cultures at recapitulating immunosenescence-mediated immune responses against cancer vaccines in vitro. The *in vivo* validation confirms these findings, as young mice demonstrated higher OVA-specific CD8+ T cell responses and smaller tumors than older mice. Our LNPoC is a valuable tool for assessing immunosenescence’s impact on cancer vaccines, potentially guiding more effective therapies for older adults.

## Introduction

Cancer vaccines are a promising type of immunotherapy that trains the immune system to recognize and attack tumor cells.^1, 2^ They work by immunizing a host against tumor-associated antigens (TAAs) to prevent tumor growth, metastasis, and recurrence.^2^ Lymph nodes (LNs) play a critical role in this immunization process, as they are the primary site for antigen presentation and T cell activation,^2, 3^ particularly within the paracortex region where the interaction between antigen-presenting cells (APCs) and T cells initiates effective antitumor immune responses. However, the clinical translation of cancer vaccines remains challenging despite their potential and global market expected to reach ∼$16 billion by 2030.^4-6^

The gap in translation can be attributed to the limitations of current preclinical models used to study cancer vaccine efficacy in recapitulating human physiology.^7, 8^ For example, the MAGE-A3 cancer vaccine showed promise in preclinical models yet failed to demonstrate clinical benefit in a Phase III trial for non-small cell lung cancer.^9, 10^ Another example is the gp100 melanoma vaccine, which initially showed potent antitumor activity in mice but failed to demonstrate similar efficacy in human clinical trials.^11^ *In vivo* murine models are the most widely used for preclinical cancer vaccine testing, but they present significant immunological differences from humans, leading to inaccurate clinical predictions of vaccine efficacy.^12, 13^ *In vitro* models, such as 2D monolayer cultures and simple 3D spheroids, also fail to replicate the complexity of the human immune system.^14^ These models do not capture the intricate spatial organization and dynamic cellular interactions within the LNs. In addition, they do not simulate *in vivo* mechanical forces and fluid dynamics, allowing the continuous flow of immune cells and antigens through lymphatic vessels. As a result, *in vitro* models fail to accurately predict the potential clinical efficacy of cancer vaccines. The limitations of current preclinical models in predicting human immune responses emphasize the need for more clinically relevant platforms for studying cancer vaccines.

With the recent passing of the FDA Modernization Act 2.0, organ-on-a-chip (OoC) technologies have gained tremendous attention for their capability to represent the human body and generate clinically translatable data.^15-17^ OoC platforms integrate microfluidic technology with cellular biology and tissue engineering to recreate the structure and function of human organs on a microscale.^18-22^ Several OoC platforms have been developed to test the efficacy of different immunotherapies;^23-29^ however, none have been developed for testing cancer vaccine efficacy. In this context, a LN paracortex-on-a-chip (LNPoC) could replicate the native microenvironment of the LN, including the critical APC-T cell interactions. Few LN-on-a-chip platforms can model the paracortical region of the lymph node where dynamic APC-T cell interactions occur, leading to T cell activation; however, none have focused on evaluating cancer vaccine responses (i.e., immunogenicity and antitumoral efficacy).^23, 30-33^ Therefore, there is a need to develop a novel physiologically relevant platform for testing cancer vaccine efficacy *in vitro*.

Another critical factor often ignored in preclinical testing of cancer vaccine efficacy is immunosenescence, the age-related decline in immune function that affects cancer patients.^34, 35^ Immunosenescence is characterized by a decrease in the population of naïve T cells, reduced T cell receptor (TCR) diversity, and an increase in the number of exhausted or senescent T cells, all of which reduce the immune system’s capacity to mount effective antitumor responses.^36-42^ This is a major hurdle for cancer vaccines, which rely on robust T cell activation and expansion to achieve therapeutic effects. Aging populations, which constitute the majority of cancer patients, are often unaccounted for in preclinical models and excluded from early-phase cancer vaccine trials,^43-45^ leading to the paucity of clinical data from these populations.^43-47^ Consequently, cancer vaccines that show efficacy in younger, healthy subjects may fail in elderly populations due to age-related immune decline or immunosenescence.^39, 48, 49^ This gap in preclinical modeling underscores the need for novel *in vitro* models that can account for the effects of immunosenescence on vaccine efficacy.

The development of a LNPoC is a boon for cancer vaccine research and development if it mimics human immunosenescence. Such technology can expedite the clinical translation of cancer vaccines, particularly for the geriatric population. Therefore, we developed a LNPoC that can recapitulate age-dependent adaptive immune responses against cancer vaccines. We conducted a proof-of-concept study to determine whether our novel LNPoC could recapitulate *in vivo* adaptive immune responses to cancer vaccines, from immunogenicity to effector functions, better than conventional 2D *in vitro* models. Using OVA-derived peptides as model antigens and murine immune cells from young or old individuals, we compared antigen presentation, antigen-specific CD8^+^ T cell generation, and antitumoral efficacy between a 2D static co-culture model, an *in vivo* B16-OVA mouse model, and our novel LNPoC.

## Experimental Methods

### Design and fabrication of the microfluidic device

The 2D computer-aided designs (CADs) for the different layers (top, middle, and bottom) of the microfluidic LNPoC were made in CorelDRAW 2021 (Corel, Canada). PMMA sheets with thicknesses of 3.18 mm (top layer) (McMaster-Carr, USA), 0.2 mm (middle layer) (Astra Products, USA), and 0.2 mm (bottom layer) (Astra Products, USA) were cut using a CO_2_ laser cutter (Universal Laser Systems, USA) based on the design shown in **Figure 1A**. Each PMMA layer was 28 mm long and 22 mm wide. The middle layer’s cut-out area consisted of a 12 mm long and 5 mm wide rounded square connected to two 3-mm diameter holes. The top layer had an inlet and outlet port (0.5 mm in diameter). After laser cutting the different PMMA layers, they were cleaned with 100% ethanol (Sigma-Aldrich, USA), rinsed with deionized water, and thoroughly dried before bonding. To bond the PMMA layers and generate the microfluidic LNPoC, the PMMA layers were treated with O_2_ plasma for 10 minutes, rinsed with 70% Isopropyl alcohol (IPA) (Sigma-Aldrich, USA), and held on top of each other for 2 minutes using paper clips. The excess IPA was removed from the microfluidic LNPoC before being placed in a 60°C oven for 15 minutes to complete the bonding of the different PMMA layers. After bonding, the resulting microchamber was washed with 70% ethanol, followed by 1X phosphate-buffered saline (PBS), and air-dried.

**Figure 1:**
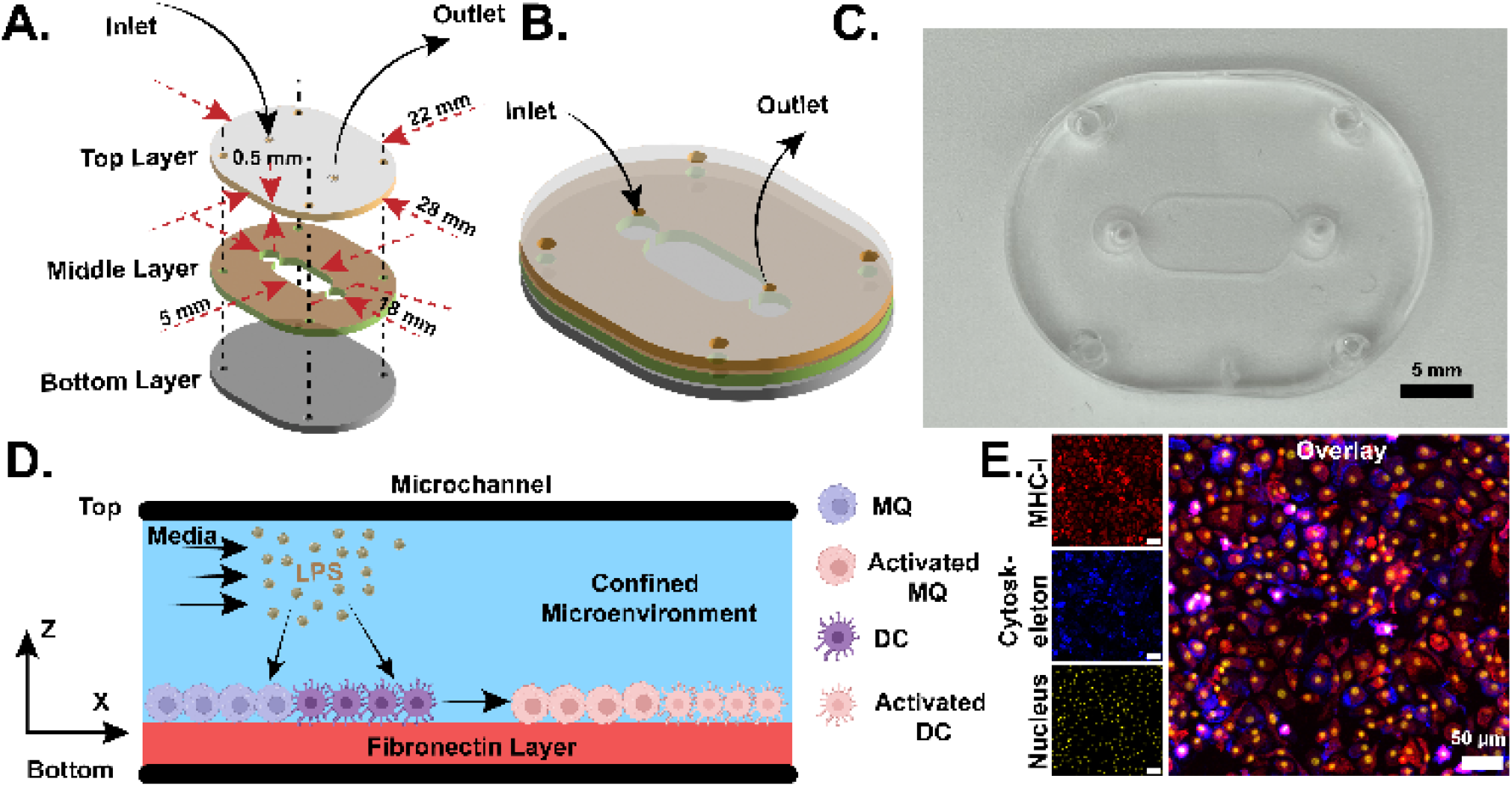
Design, fabrication, and characterization of the device. **A.** Design of the LNPoC. **B.** LNPoC after assembly. **C.** Optical image of the LNPoC device. **D.** Schematic diagram of the device with coating and attached cells. **E.** APCs attached inside the LNPoC device.

### Isolation of monocytes from mice bone marrow

Animals received comprehensive care in compliance with federal, state, and local guidelines. All procedures conducted on animals adhered to the regulations and were approved by the Lundquist Institute Committee on Use and Care of Animals (ICUCA #32706-02). Young (6-7 weeks old) and old (35-36 weeks old) C57BL/6 female mice (The Jackson Laboratory, USA) were housed in a pathogen-free environment for at least 3 days and kept at room temperature with free access to water and a 12-hour light/dark cycle. C57BL/6 mice were euthanized, and the femurs and tibia were harvested. The bone marrow was flushed out of the femurs and tibia with sterile 1X PBS using a 5 mL syringe with a 30-gauge needle, and the bone marrow was collected on a 40 µm cell strainer (Corning, USA) placed on top of a 50 mL conical tube. The bone marrow was mashed using a syringe’s plunger, and then the cell strainer was rinsed with 10 mL of 1X PBS. The resulting bone marrow cell suspension was centrifuged at 400 x g for 10 minutes at 4°C. The cells were resuspended in 1X PBS and directly used for subsequent differentiation into MQs and DCs.

### Macrophages and dendritic cell differentiation

The freshly isolated young or old bone marrow cells were plated in a Petri dish (2 x 10^6^ cells/dish) and cultured for 7 days at 37°C with 5% CO_2_, and half of the culture media was replaced with fresh media on days 3 and 5. To generate MQs, the bone marrow cells were cultured in Dulbecco’s Modified Eagle’s Medium (DMEM) + GlutaMax (Fisher Scientific, USA), 1X non-essential amino acid (Thermo Fisher Scientific, USA), 1 mM sodium pyruvate (Cytiva, USA), 100 µM B-Mercaptoethanol (Sigma-Aldrich, USA), and mouse macrophage colony-stimulating factor (M-CSF) (5ng/mL) (Miltenyi Biotec, USA). To generate DCs, the bone marrow cells were cultured in Roswell Park Memorial Institute (RPMI) (Cytiva, USA), 1X non-essential amino acid, 1 mM sodium pyruvate, 100 µM B-Mercaptoethanol, and mouse granulocyte-macrophage colony-stimulating factor (GM-CSF) (5ng/mL) (GenScript, USA). The differentiation of bone marrow cells into MQs (CD11b^+^) or DCs (CD11c^+^) was confirmed with flow cytometry (FACSCanto II flow cytometer, BD Biosciences, USA). Using FlowJo^TM^ software (Becton, Dickinson and Company, USA), Differentiated bone marrow cells were first gated based on size (side-scatter) and granularity (forward-scatter), then MQs (F4/80^+^/CD11b^+^) were quantified after staining the cells with an Alexa Fluor® 700-conjugated monoclonal anti-mouse F4/80 Antibody (BioLegend, USA) and a Brilliant Violet 711^TM^-conjugated monoclonal anti-mouse CD11b antibody (BioLegend, USA), and DCs (F4/80^-^/CD11c^+^) were quantified after staining the cells with a PerCP-conjugated monoclonal anti-mouse F4/80 Antibody (BioLegend, USA) and a FITC-conjugated monoclonal anti-mouse CD11c antibody (BioLegend, USA).^50, 51^

### Stimulation of macrophages and dendritic cells using OVA peptide

Young and old MQs and DCs were collected from Petri dishes and resuspended at a concentration of 1×10^6^ cells/mL in their corresponding cell culture media containing 100 ng/mL of OVA peptide fragment (SIINFEKL) (Anaspec, USA) and 250 ng/mL of lipopolysaccharide (LPS) (Sigma-Aldrich, USA). For the off-chip stimulation, 100 µL of MQs or DCs suspensions were seeded in a flat-bottom 96-well plate and cultured for 7 days at 37°C with 5% CO_2_, and half of the culture media was replaced with fresh media on days 3 and 5. For the on-chip stimulation, the LNPoC was first sterilized with 30 minutes of UV treatment, followed by 70% ethanol wash for 10 minutes. The microfluidic devices were washed 5 times with sterile 1X PBS, then coated with fibronectin (50 µg/mL) (Thermo Fisher Scientific, USA) for 2 hours at 37°C and 5% CO_2_. Finally, 20 µL of young or old cell suspension were seeded and cultured in the LNPoC for 7 days at 37°C with 5% CO_2_, and half of the culture media was replaced with fresh media on days 3 and 5. The presentation of the OVA peptide by MHC-I on MQs or DCs was analyzed by flow cytometry using FlowJo^TM^ software. Differentiated bone marrow cells were first gated based on size (side-scatter) and granularity (forward-scatter), then MQs (OVA^+^/CD11b^+^) and DCs (OVA^+^/CD11c^+^) were quantified after staining the cells with an iTAg Tetramer/PE-H-2 Kb OVA (NIH Core Facility, USA), a Brilliant Violet 711^TM^-conjugated monoclonal anti-mouse CD11b antibody (BioLegend, USA), and a FITC-conjugated monoclonal anti-mouse CD11c antibody (BioLegend, USA).

### Isolation of naïve CD8^+^ T cells from mice spleen

Young (6-7 weeks old) and old (35-36 weeks old) C57BL/6 female mice (The Jackson Laboratory, USA), housed in a pathogen-free environment for at least 3 days and kept at room temperature with free access to water and a 12-hour light/dark cycle, were euthanized under the ICUCA committee of the Lundquist Institute (#32706-02), and spleens were harvested. Under sterile conditions, four spleens were cut into small pieces on a petri dish using a scalpel and then soaked in T cell media (RPMI, 1X non-essential amino acid, 1 mM sodium pyruvate, and 100 µM B-Mercaptoethanol). The solution containing the young or old minced spleens was passed through a 40 µm cell strainer placed on top of a 50 mL conical tube. The spleens were mashed using a syringe’s plunger, and then the cell strainer was rinsed with 10 mL of T cell media. The resulting spleen cell suspension was centrifuged at 400xg for 10 minutes at 4°C, and the cell pellet was resuspended in T cell media. Then, the spleen cell suspension was passed through a 40 µm cell strainer to remove clumps or debris. The young and old naïve CD8^+^ T cells were isolated by negative selection from the purified spleen cell solution using the EasySep™ Mouse CD8^+^ T Cell Isolation Kit (STEMCELL Technologies, USA). The purification of naïve CD8^+^ T cells was confirmed by flow cytometry using FCS Express^TM^ software. Purified cells were first gated based on size (side-scatter) and granularity (forward-scatter), then live naïve CD8^+^ T cells were quantified after staining the cells with a Zombie Violet™ Fixable Viability Kit (BioLegend, USA) and an APC-conjugated recombinant anti-mouse CD8a antibody (BioLegend, USA).

### OVA-specific activation of naïve CD8^+^ T cells

Freshly isolated young or old naïve CD8^+^ T cells were resuspended in T cell media and then co-cultured with OVA-stimulated MQs and DCs. For the off-chip activation of naïve CD8^+^ T cells, 100 µL of MQs and DCs suspension (1:1 MQs to DCs ratio at a concentration of 1×10^6^ cells/mL) were seeded in a flat-bottom 96-well plate and cultured in cell culture media containing 100 ng/mL of OVA peptide and 250 ng/mL of LPS for 7 days at 37°C with 5% CO_2_, and half of the culture media was replaced with fresh media on days 3 and 5. After 7 days of OVA peptide stimulation, 100 µL of naïve CD8^+^ T cells (1×10^6^ cells/mL) were added to the 96-well plate and co-cultured for 2 days at 37°C and 5% CO_2_ with MQs and DCs. After 2 days, activated CD8^+^ T cells were collected from the 96-well plate. For the on-chip activation of naïve CD8^+^ T cells, 20 µL of young and old MQs and DCs suspension were seeded and cultured in the LNPoC with cell culture media containing 100 ng/mL of OVA peptide and 250 ng/mL of LPS for 7 days at 37°C with 5% CO_2_, and half of the culture media was replaced with fresh media on days 3 and 5. After 7 days, 20 µL of naïve CD8^+^ T cells were seeded into the LNPoC containing OVA-stimulated MQs and DCs and cultured for 2 days at 37°C and 5% CO_2_. After 2 days, activated CD8^+^ T cells were collected from the LNPoC. The generation and proliferation of young and old OVA-specific CD8^+^ T cells were analyzed by flow cytometry using FlowJo^TM^ software. Activated CD8^+^ T cells were first gated based on size (side-scatter) and granularity (forward-scatter), then OVA-specific CD8^+^ T cells were quantified after staining the cells with a PE-conjugated OVA-tetramer (NIH core facility, USA) and an APC-conjugated recombinant anti-mouse CD8a antibody (BioLegend, USA). Activated CD8^+^ T cells were also stained with CFSE (Thermo Fisher Scientific, USA) to assess the proliferation of OVA-specific CD8^+^ T cells.

### Simulation of velocity and shear stress distribution in the device

A 3D computational model was developed in COMSOL Multiphysics using the finite element-based method to simulate media flow inside the device (**Figure S1**). A single-phase, laminar flow model was used, with the fluid (water) assumed to be isothermal, incompressible, and Newtonian. The governing equations for this simulation were as follows ^52^:

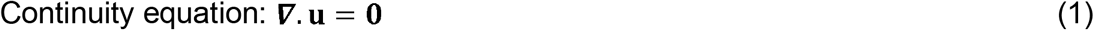

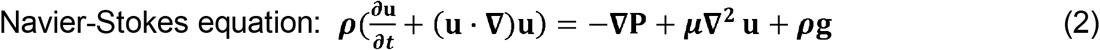

The symbols **u, *ρ*, *µ*, g,** and **P** refer to velocity, the density of the water, dynamic viscosity, acceleration due to gravity, and pressure, respectively. The parameters of the simulations are as follows: u= 0.00017 m/s, **u**_i_ (at t = 0 s) = 0, ***µ*** = 0.00089 Pa s, g = O, and ***ρ*** = 1000 kg/m^3 53-56^. The boundaries of the simulated domain were selected to be impermeable and non-slipping (**u**= 0). The computational domain was discretized into 124,943 triangular elements to ensure a mesh-independent solution. The spatiotemporal distributions of velocity and wall shear stress were obtained and visualized. A cutline (X = 8 mm, Y = 2.5 mm, and Z = 0 to 0.2 mm) was placed across the channel height to extract one-dimensional velocity and shear stress profiles (**Figure S1**).

### Antitumoral assay

The B16-OVA cells (Sigma-Aldrich, USA) were cultured in DMEM (Fisher Scientific, USA), supplemented with 1% penicillin-streptomycin (Caisson Labs, USA) and 10% fetal bovine serum (FBS) (Life Technologies, USA). The B16-OVA cells were seeded in 6-well plates and grown until confluency (≈1×10^6^ cells/well). Then, the young and old activated CD8^+^ T cells collected from the LNPoC were added to each well (1×10^6^ cells/well) and co-cultured with B16-OVA cells. After 2 days, CD8^+^ T cells and B16-OVA cells are collected and analyzed by flow cytometry using FCS Express^TM^ software. The cells were first gated based on size (side-scatter) and granularity (forward-scatter), and the viability of B16-OVA cells was quantified after staining the cells with a Zombie Violet™ Fixable Viability Kit (BioLegend, USA) and an APC-conjugated recombinant anti-mouse CD8a antibody (BioLegend, USA).

### *In vivo* animal study

Animals received comprehensive care in compliance with federal, state, and local guidelines. All procedures conducted on animals adhered to the regulations and were approved by the Lundquist Institute Committee on Use and Care of Animals (ICUCA #32706-02). Young (6-7 weeks old) and old (35-36 weeks old) C57BL/6 female mice (The Jackson Laboratory, USA) were housed in a pathogen-free environment for at least 3 days and were kept at room temperature with free access to water and a 12-hour light/dark cycle. The *in vivo* study involved C57BL/6 female mice bearing B16-OVA tumors. B16-OVA cells, cultured until 80% confluence and trypsinized with TripleE (Fisher Scientific, USA), were injected (1×10^6^ cells in 100 µL /mouse) subcutaneously into the right flank of young and old C57BL/6 mice. After cell injection, the growth of B16-OVA tumors was monitored every other day. Tumor size was measured using a caliper and recorded, and the tumor volume was calculated using the formula: tumor volume (mm^3^) = length × (width)^2^ × 0.5. Once the tumors reached 20 mm^3^, the young and old mice were subcutaneously immunized at the tail base on days 0, 5, and 10 with 100 µL of free OVA peptide (1 mg/mL) or PBS (negative control). On days 7, 14, and 21 post-first immunization, mice were anesthetized, and blood samples (100 μl) were collected submandibularly. Animals were euthanized by CO_2_ asphyxiation when they reached either of the following end-points: (1) day 21 post-first immunization; (2) the tumor mass reached 1.5 cm in any dimension (length or width); or (3) animals exhibited moribund conditions (>20% weight loss or ulceration). Once euthanized, tumor and LN tissues were harvested and processed. OVA-specific CD8^+^ T cells in tumors, lymph nodes, and blood samples were quantified by flow cytometry using FCS Express^TM^ software. CD8^+^ T cells were first gated based on size (side-scatter) and granularity (forward-scatter), then OVA-specific CD8^+^ T cells were quantified after staining the cells with a PE-conjugated OVA-tetramer (NIH core facility, USA) and an APC-conjugated recombinant anti-mouse CD8a antibody (BioLegend, USA).

### ELISA

The levels of cytokines in the media collected after 7 days of MQs and DCs stimulation using OVA peptide were quantified using IL-6, IL-10, IL-12, TNF-alpha, and IFN-gamma ELISA kits (BioLegend, USA) following the manufacturer’s protocol. The absorption measurements used to quantify the concentrations of IL-2, IL-6, IL-10, and IL-12 were acquired with the Varioskan™ LUX multimode microplate reader (Thermo Fisher Scientific, USA).

### Statistical analysis

All statistical analyses were performed using the GraphPad Prism 9 software. Results are expressed as mean ± standard deviation from triplicate experiments (N = 3). All data was tested for normality using the Shapiro-Wilk and Shapiro-Francia tests. When applicable, analyses for significance were performed using a parametric or non-parametric analysis of variance (ANOVA) test. Statistical significance was defined by p<0.05 for two-sided p-values.

## Results

### Microfluidic chip fabrication and characterization

We fabricated a microfluidic LNPoC device using Poly(methyl methacrylate) (PMMA). The device comprised a rounded square cell culture microchamber (length: 12mm; and width: 5mm) connected to two 3 mm diameter holes leading to the inlet and outlet ports (diameter: 0.5mm) (**Figure 1A-C**). Once fabricated, the LNPoC device was coated with fibronectin, which promoted macrophages (MQs) and dendritic cells (DCs) adhesion on the bottom of the cell culture microchamber and allowed the culture of MQs and DCs within the device (**Figure 1D**). This fibronectin coating formed a confined microenvironment appropriate for the cell culture, activation, and antigen presentation by APCs inside the LNPoC device. Successful attachment and distribution of APCs within the device were confirmed through immunofluorescence imaging of MHC-I, cytoskeleton, and nuclei staining (**Figure 1E**).

### LNPoC revealed age-dependent presentation of OVA peptide

To evaluate the ability of the LNPoC platform to model age-related differences in antigen presentation, we isolated monocytes from the bone marrow of young and old mice and differentiated them into MQs and DCs. A schematic representation of the experimental workflow depicts the process of isolating monocytes, differentiating them into MQs and DCs, and loading the OVA peptide either on-chip or off-chip to evaluate antigen presentation capacity (**Figure 2A**). We isolated bone marrow cells from the tibia and femur of young (6–7 weeks old) and old (35–36 weeks old) C57BL/6 mice. Prior to initiating the differentiation protocol, we confirmed that the starting cell populations were devoid of macrophages (F4/80^+^/CD11b^+^) and dendritic cells (F4/80^?^/CD11c^+^), ensuring that only undifferentiated monocytes were present for subsequent experiments (**Figure 2B**).^57, 58^ After 7 days of differentiation, we confirmed for young (**Figure 2C**) and old (**Figure 2D**) mice that bone marrow cells were differentiated into MQs (F4/80^+^/CD11b^+^) and DCs (F4/80^-^/CD11c^+^). The flow cytometry plots demonstrate that immune cells derived from young mice exhibit higher activation and antigen presentation levels than those from older mice. Specifically, young mouse-derived macrophages and dendritic cells show increased expression of activation markers and a greater capacity to present OVA peptides. This enhanced antigen presentation leads to more robust activation of OVA-specific CD8^+^ T cells, critical for effective antitumor immune responses. These findings highlight the age-dependent differences in immune function and underscore the importance of using physiologically relevant models like the LNPoC to study immunosenescence and its impact on cancer vaccine efficacy.

**Figure 2:**
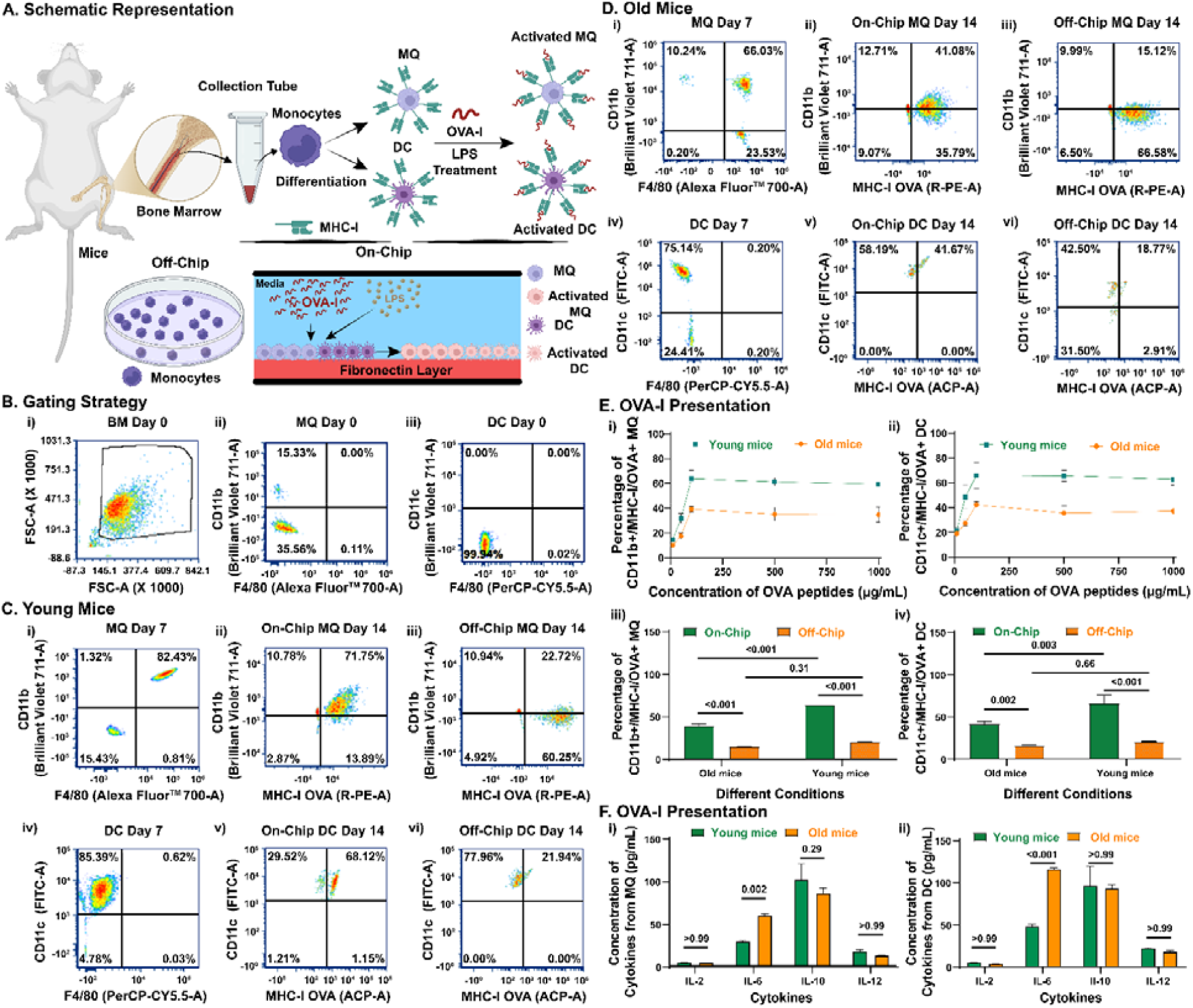
Differentiation of mouse bone marrow cells into macrophages (MQs) and dendritic cells (DCs), OVA peptide presentation, and cytokine secretion. **A.** Schematic representation of the differentiation of monocytes isolated from mouse bone marrow into MQs and DCs, followed by OVA peptide loading either on-chip or off-chip. Representative graphs showing dot plots from flow cytometry analysis: **B.** After isolation of (i) bone marrow cells from young and old mice, confirming the absence of (ii) MQs and (iii) DCs before differentiation. After 7 days of differentiation on-chip and off-chip in **C.** young and **D.** old murine immune cells with OVA presentation on day 14: (i) differentiated MQs at Day 7, (ii) on-chip OVA presented MQs on Day 14, (iii) off-chip OVA presented MQs on Day 14, (i) differentiated DCs at Day 7, (ii) on-chip OVA presented DCs on Day 14, (iii) off-chip OVA presented DCs on Day 14. **E.** Dose-dependent OVA presentation and percentage of OVA presentation on MQs and DCs: (i) dose-dependent OVA presentation on MQs, (ii) dose-dependent OVA presentation on DCs, (iii) percentage of OVA presentation on MQs and (iv) percentage of OVA presentation on DCs. **F.** On-chip cytokine secretion at day 14 for old and young mice: from (i) MQs and (ii) DCs.

After 7 more days of stimulation with OVA peptide, we analyzed the OVA peptide presentation potential of MQs (OVA^+^/CD11b^+^) or DCs (OVA^+^/CD11c^+^) in young and old cells. We observed that OVA peptide was presented by young and old MQs and DCs, both on-chip and off-chip. On-chip, we observed a dose-dependent OVA peptide presentation response and a plateau of presentation when a 100 μg/mL concentration of OVA peptide was used as the immunogen (**Figure 2E**). This dosage was used subsequently for all experiments and analyses. For the OVA peptide presentation by MQs, we observed a significant difference in OVA peptide presentation between on-chip and off-chip experiments using cells from young (63.8 ± 4.9% vs. 20.7 ± 0.2%, p<0.001) and old mice (39.3 ± 2.6% vs. 15.2 ± 0.2%, p<0.001). In addition, we observed that on-chip OVA peptide presentation was significantly higher with young cells than old ones (63.8 ± 4.9% vs. 39.3 ± 2.6%, p<0.001), whereas no significant difference in off-chip OVA peptide presentation between young cells and old ones (20.7 ± 0.2% vs. 15.2 ± 0.2%, p=0.31). The same observations were made for DCs when comparing on-chip and off-chip results using young (65.9 ± 10.1% vs. 21.2 ± 0.5%, p<0.001) and old cells (42.3 ± 2.0% vs. 16.1 ± 0.9%, p=0.002), and on-chip OVA peptide presentation with young cells (65.9 ± 10.1% vs. 42.3 ± 2.0%, p=0.003) and old ones (21.2 ± 0.5% vs. 16.1 ± 0.9%, p=0.66) (**Figure 2E**). The confocal images of the on-chip presentation of the OVA peptide confirmed that the OVA presentation by young MQs and DCs was higher than that of old MQs and DCs (**Figure S2**). Lastly, we assessed the cytokine production from MQs and DCs on day 7 after on-chip OVA peptide and lipopolysaccharide (LPS) stimulation, and we observed a significant difference in IL-6 secretion between young and old MQs (p=0.002) and DCs (p<0.001). We did not observe a significant difference for IL-2, IL-10, and IL-12 (**Figure 2F**).

### LNPoC revealed the age-dependent generation of OVA-specific CD8^+^ T cells

To investigate how aging affects the activation and proliferation of antigen-specific T cells, we isolated CD8^+^ T cells from the spleens of young and old C57BL/6 mice. We then analyzed their functional responses using the LNPoC platform. A schematic representation of the experimental workflow illustrates the process of isolating CD8^+^ T cells from mouse splenocytes, co-culturing them with OVA peptide-presenting MQs and DCs within the LNPoC and following the subsequent activation and proliferation of OVA-specific CD8^+^ T cells in a controlled microenvironment (**Figure 3A**). We isolated CD8^+^ T cells through a negative selection process, resulting in a cell fraction containing 34.8% and 49.5% of live CD8^+^ T cells from young and old mice, respectively (**Figure 3B**). Next, we characterized OVA-specific T cell proliferation and OVA peptide specificity in young (**Figure 3C**) and old (**Figure 3D**) immune cells. We observed that using young and old immune cells, the proliferation of T cells was significantly higher on-chip than off-chip (p<0.001).

**Figure 3:**
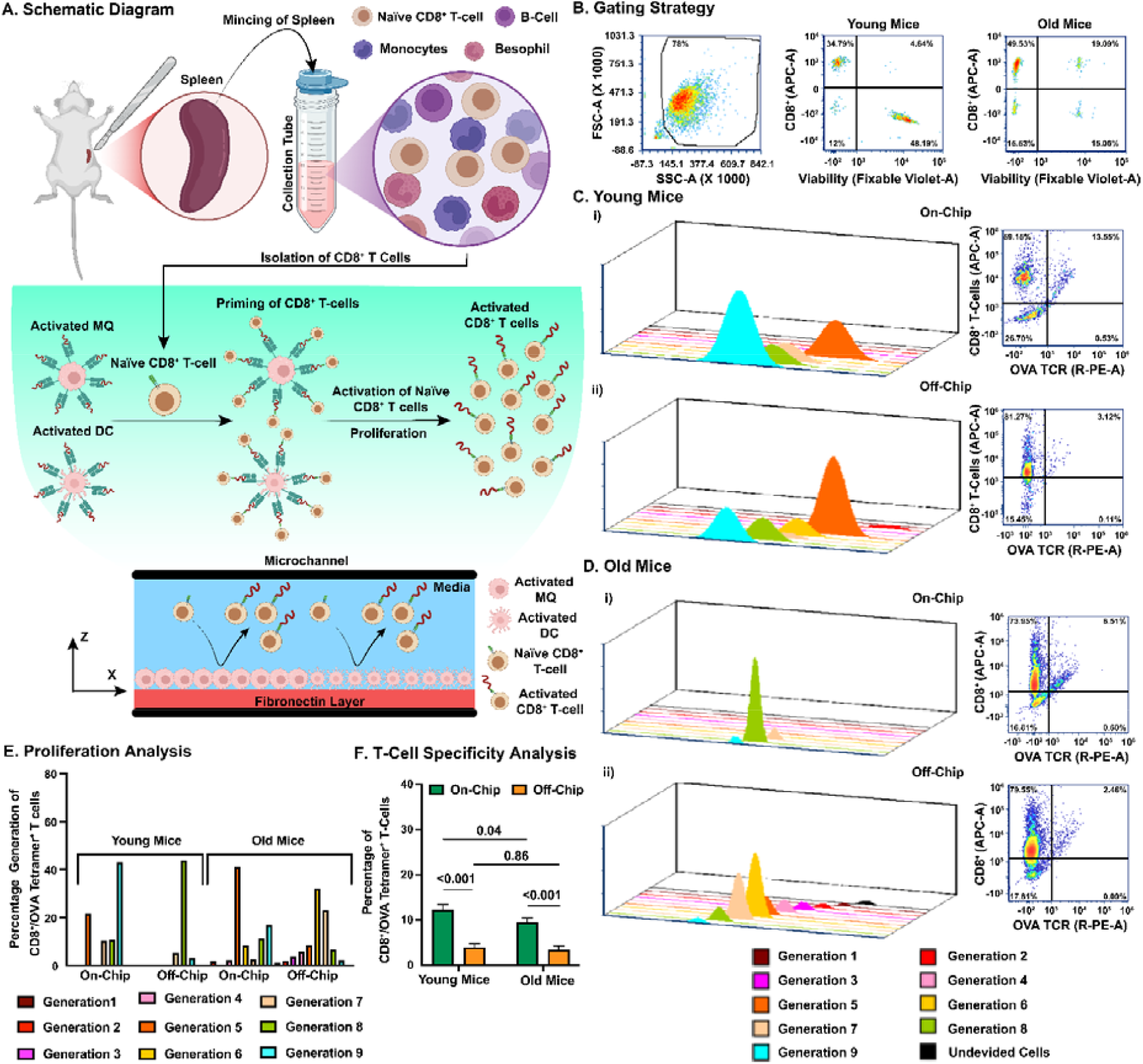
OVA-I-specific CD8+ T cell activation. **A.** Schematic diagram of the activation of CD8+ T-cells on-chip. **B.** Total population of splenocytes, CD8+ T cells from young and old mice were gated based on forward-and side-scatter profiles: i) Isolated splenocytes; ii) Negative selection of CD8+ T cells from splenocytes from young mice; iii) Negative selection of CD8+ T cells from splenocytes from old mice. **C.** On-chip and off-chip OVA-specific CD8+ T cell proliferation using CFSE (i) and CD8+ T cell activation (ii) after 3 days of co-culture with OVA-I presenting MQs and DCs from young mice. **D.** On-chip and off-chip OVA-specific CD8+ T cell proliferation using CFSE (i) and CD8+ T cell activation (ii) after 3 days of co-culture with OVA-I presenting MQs and DCs from old mice. **E.** Comparison of OVA-I-specific CD8+ T cell generation on-chip and off-chip in young and old mice. **F.** Comparison of percentage of OVA-I-specific CD8+ T cells on-chip and off-chip in young and old mice.

Also, we observed a significantly higher on-chip proliferation of young CD8^+^ T cells than old CD8^+^ T cells (p<0.001); however, this difference was not observed off-chip (**Figure 3E**). Generations 1-9 demonstrate the proliferation of OVA-specific CD8^+^ T cells, measured by CFSE staining. Young CD8^+^ T cells show more cells in the later generations (5-9), indicating stronger proliferation. In contrast, old CD8^+^ T cells mostly remain in the earlier generations (1-4), suggesting impaired proliferation due to immunosenescence. In a similar way, we observed a significantly higher generation of OVA-specific CD8^+^ T cells on-chip compared to off-chip for young (12.3 ± 1.3% vs. 4.0 ± 0.8%, p<0.001) and old (9.5 ± 0.9% vs. 3.4 ± 1.0%, p<0.001) CD8^+^ T cells (**Figure 3F**). We also observed a significantly higher percentage of OVA-specific T cells on-chip using young T cells compared to old ones (12.3 ± 1.3% vs. 9.5 ± 0.9%, p=0.04), whereas no significant differences were observed between young and old cells (4.00 ± 0.8% vs. 3.36 ± 1.0%, p=0.86) (**Figure 3F**). Confocal images of on-chip activation of OVA-specific CD8^+^ T cells confirmed higher OVA priming of young CD8^+^ T cells than old CD8^+^ T cells (**Figure S3**).

To isolate the activated CD8^+^ T cells within the device, we employed flow-induced shear stress. We varied the media velocity to optimize the cell isolation process. The spatiotemporal distribution of the isolated CD8^+^ T cells was assessed within the LNPoC over a time span of 0 to 30 seconds, confirming the minimal optimal velocity (average velocity: 0.00017 m/s) necessary to separate the activated cells from the total population of CD8^+^ T cells (**Figure S1A**). We also conducted a corresponding CFD simulation to model the velocity and shear stress distributions. The geometry of the LNPoC device used for the CFD simulations consists of a 10 mm × 5 mm microchannel with a height of 0.2 mm and circular inlet and outlet ports (**Figure S1B**). The CFD simulations indicated the spatial velocity (**Figure S1C**) and shear stress (**Figure S1D**) distributions across the device at an average velocity of 0.00017 m/s, highlighting a uniform flow along the microchannel. The velocity profile across the channel height exhibited a parabolic distribution characteristic of laminar flow, with peak velocity increasing proportionally to the inlet velocity (**Figure S1E**). Similarly, the shear stress profile was highest near the walls and increased with the inlet velocity, ranging from 0.001 to 0.008 Pa (**Figure S1E**). These results validate the controlled microenvironment and physiologically relevant flow conditions within the LNPoC.

### Antitumoral efficacy of OVA-I-specific CD8^+^ T cells

We assessed the antitumoral efficacy of CD8^+^ T cells stimulated by OVA-presenting MQs and DCs by co-culturing B16-OVA cells with activated CD8^+^ T cells. A schematic illustration of the experimental workflow displays the co-culture of activated CD8^+^ T cells with B16-OVA melanoma cells and subsequent assessment of tumor cell death (**Figure 4A**). To isolate CD8^+^ T cells, we used a negative selection process, which removes unwanted blood cell types while leaving CD8^+^ T cells untouched. This method relies on depleting other immune cells, such as B cells, NK cells, and myeloid cells, resulting in an enriched population of CD8^+^ T cells. Additionally, our analysis included gating on all live cells rather than exclusively on CD8^+^ T cells. Because negative selection eliminates many non-T cell populations, the remaining cell suspension contains a higher proportion of CD8^+^ T cells among all viable cells. This explains why we observed 34.8% live CD8^+^ T cells from young mice and an even higher 49.5% from old mice. The increased percentage in older mice may also reflect age-related shifts in immune cell composition. Still, the main factor is the combination of negative selection and the broad gating strategy, which measures CD8^+^ T cells as a fraction of all remaining live cells rather than just among T cells. Compared to a negative control without OVA-specific CD8^+^ T cells (**Figure 4B**), we assessed the antitumoral efficacy of young and old CD8^+^ T cells generated on-and off-chip (**Figure 4C-F**). We observed a significantly higher percentage of dead B16-OVA cells when OVA-specific CD8^+^ T cells were generated on-chip compared to off-chip for young (49.9 ± 1.5% vs. 15.1 ± 0.7%, p<0.001) and old (32.7 ± 4.0% vs. 7.0 ± 0.9%, p<0.001) immune cells (**Figure 4G**). Similarly, we observed a significantly higher percentage of dead B16-OVA cells with on-chip-generated young CD8^+^ T cells compared to old ones (49.9 ± 1.5% vs. 32.7 ± 4.0%, p<0.001), and the same trend was observed for OVA-specific CD8^+^ T cells generated off-chip (15.1 ± 0.7% vs. 7.0 ± 0.9%, p=0.01). Confocal images also confirmed more dead B16-OVA cells with young OVA-specific CD8^+^ T cells than old ones (**Figure S4**).

**Figure 4:**
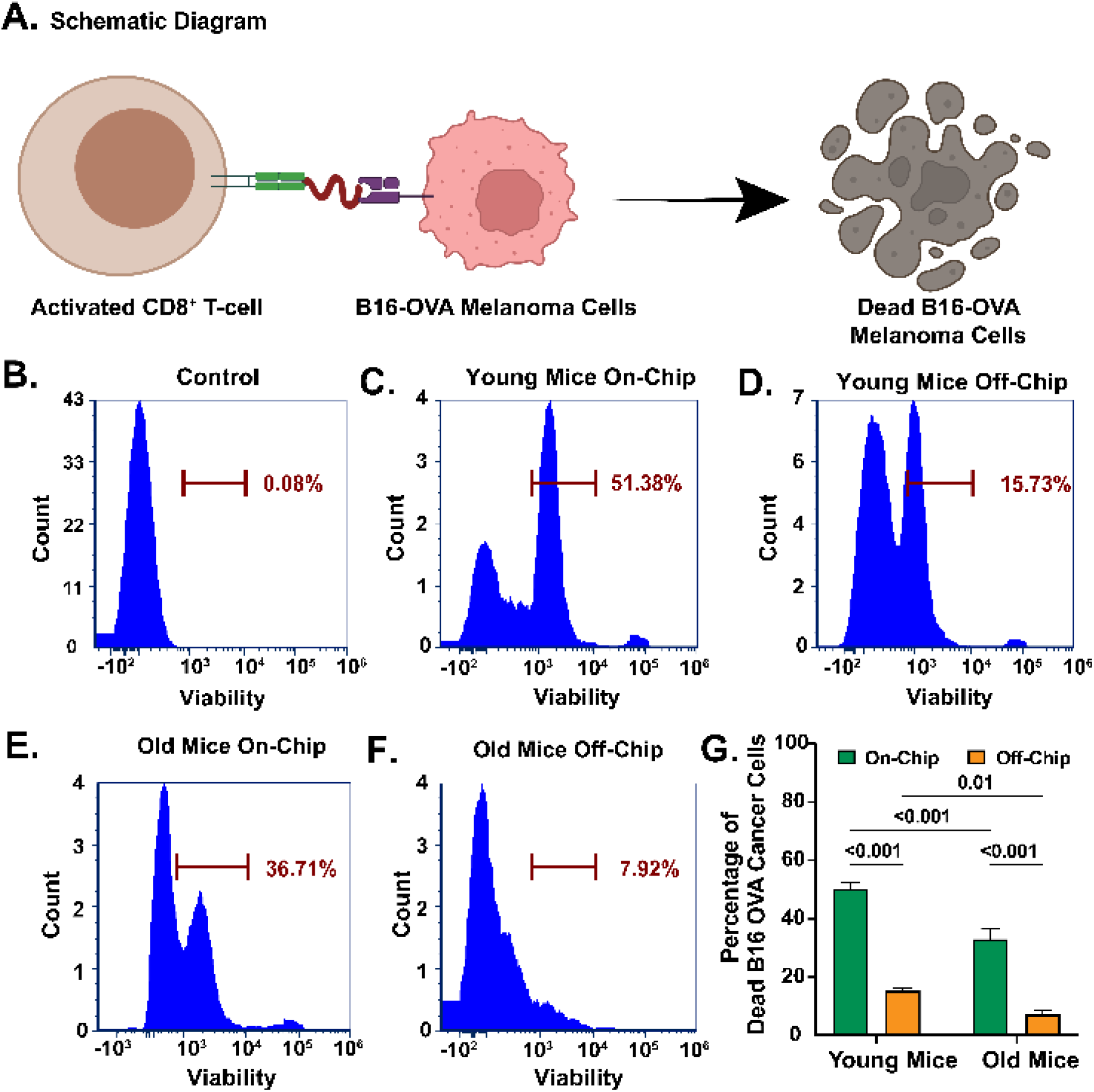
Antitumoral efficacy of OVA-I-specific CD8+ T-cells enriched in the cancer vaccine-on-a-chip platform. **A.** The schematic of the antitumoral efficacy of CD8+ T cells against B16-OVA. **B.** Negative control without OVA-specific CD8+ T-cells. **C-F.** Antitumoral efficacy of enriched OVA-I-specific CD8+ T-cells against B16-OVA melanoma cells between young and old mice and on- and off-chip with control. **G**. Percentage of dead B16-OVA melanoma cells in young and old mice and on- and off-chip.

### *In vivo* validation of the LNPoC platform

To validate the results from our *in vitro* platform, we evaluated the immunogenicity and antitumoral efficacy of OVA peptides (without adjuvants) in a subcutaneous B16-OVA murine melanoma model. We inoculated the B16-OVA cells and collected blood on day 0. We immunized the mice with OVA peptides and collected blood one week after inoculation. We also collected blood after two weeks. We sacrificed the mice after three weeks and collected blood, LNs, and B16-OVA cells (**Figure 5A**). Although not significant, the tumor growth was lower in young mice compared to old ones (**Figure 5B**). The percentage of circulating OVA-specific CD8^+^ T cells is significantly higher in PBMC of young mice compared to old mice on day 5 (11.4 ± 1.4% vs. 7.1 ± 1.0%, p<0.001), 14 (12.8 ± 1.3% vs. 8.2 ± 0.8%, p<0.001), and 21 (12.4 ± 0.8% vs. 8.8 ± 0.9%, p<0.001) (**Figure 5C**). Similarly, a significantly higher percentage of lymph node-isolated (13.4 ± 0.9% vs. 7.7 ± 2.0%, p<0.001) and tumor-infiltrating (27.3 ± 2.5% vs. 13.6 ± 1.2%, p<0.001) OVA-specific CD8^+^ T cells is in young mice compared to old mice was observed on day 21 (**Figure 5D-E**). Notably, it was observed that the percentages of OVA-specific CD8^+^ T cells in young mice present in the circulation and isolated from LNs at day 21 were similar to the on-chip young OVA-specific CD8^+^ T cells (**Table S1**).

**Figure 5:**
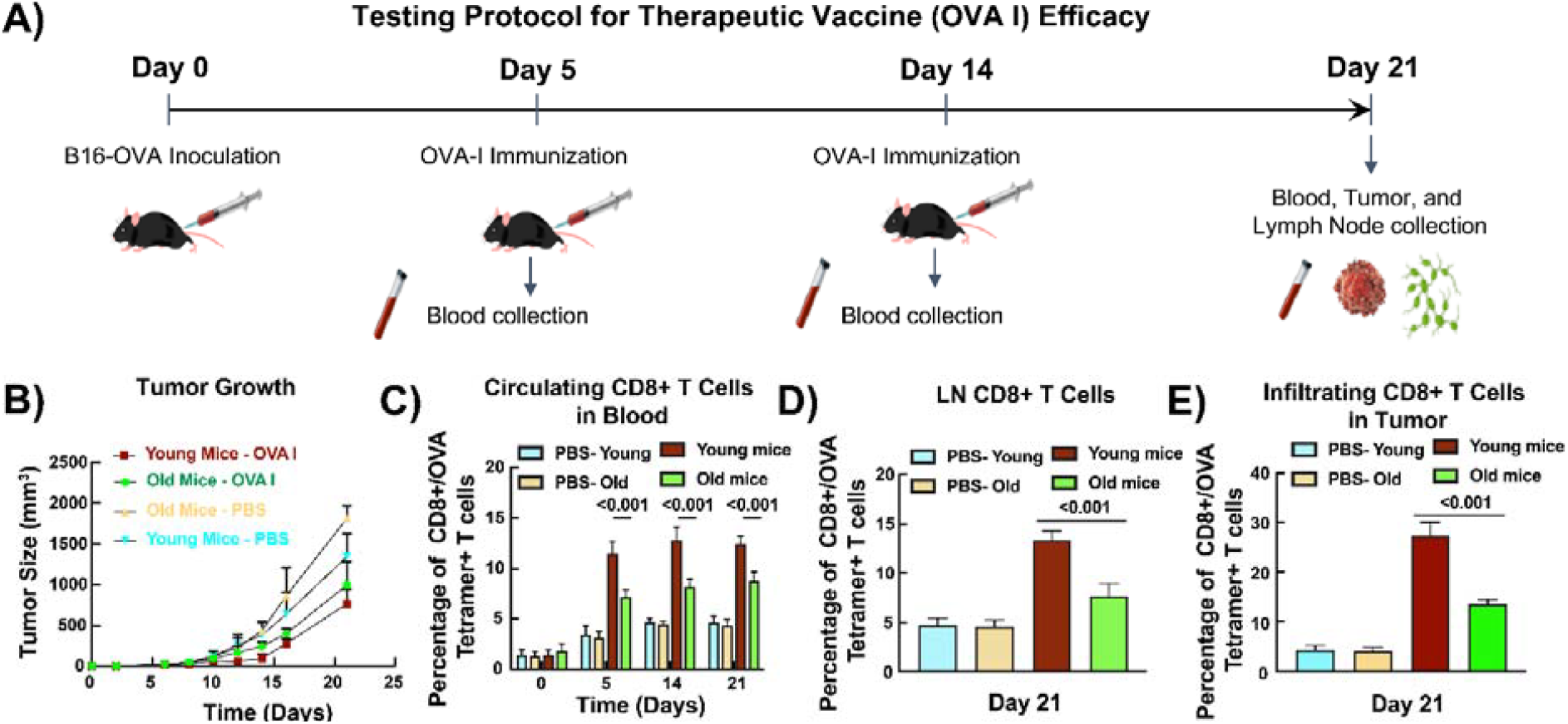
In vivo validation of the cancer vaccine-on-a-chip platform. **A.** Protocol for testing the immunogenicity and antitumoral efficacy of the OVA-I peptide vaccine against the B16-OVA flank mouse model. **B.** Tumor volume. **C.** Percentage of tetramer-positive T cell in PBMC. **D.** Percentage of tetramer-positive T cells in the LNs. **E.** Percentage of tetramer-positive tumor infiltrating CD8+ T cells.

## Discussion

We developed a LNPoC capable of recapitulating *in vivo* cancer vaccine immunogenicity in an age-dependent manner, including critical functions of the LN paracortex, such as antigen presentation and antigen-specific T cell activation, proliferation, and effector functions. Our LNPoC platform permitted the flow, capture, processing, and presentation of OVA peptides (as a model antigen) and the migration of T cells over APCs, promoting the interactions necessary for eliciting vaccine-induced immune responses. These functions are integral to mounting an effective antitumor immune response and are known to be compromised with age.^36, 59^ In contrast to conventional 2D co-culture systems, which fail to capture the spatial complexity of lymphoid tissue, the LNPoC demonstrated age-dependent differences in immune functions, demonstrating its potential as a superior model for studying immunosenescence *in vitro*.^60, 61^ This novel platform provides a robust model to examine immune responses in an age-dependent manner, where older immune cells may be less capable of initiating strong antitumor responses following cancer vaccine immunization than younger cells.

The LNPoC platform demonstrated distinct age-related differences in antigen presentation, which were not revealed off-chip using conventional 2D co-culture systems. By comparing immune responses between young and old immune cells, we demonstrated that APCs (MQs and DCs) from young mice presented OVA peptides more effectively than those from older mice on-chip but not off-chip. This significant difference in antigen presentation is consistent with previous findings that aging leads to diminished functions and impaired antigen presentation in MQs and DCs.^38, 62^ Immunosenescent DCs display reduced phagocytosis and the cross-presentation of the cell-associated antigens due to the increased oxidative stress associated with aging.^62-64^ Similarly, MQs have shown a decrease in the ability to activate the Toll-like receptors (TLR), which play a critical role in innate immune responses against cancer vaccines.^65-67^ Indeed, aging impacts cytokine secretion by APCs, especially on IL-6.^68^ IL-6 plays an essential role in pro-inflammatory immune responses and age-related chronic inflammations, resulting in altered signaling pathways, increased production of reactive oxygen species, and chronic low-grade inflammation.^69-76^ Our study confirmed the upregulation of IL-6 in old APCs compared to young ones. The decline in OVA peptide presentation by MQs and DCs from old mice is consistent with established features of immunosenescence, including impaired antigen processing, reduced TLR activation, and altered cytokine secretion profiles in aged APCs.^77, 78^

We also demonstrated a significant reduction in the generation, proliferation, and activation of OVA-specific CD8^+^ T cells in old cells compared to young ones. These results align with established evidence that aging is associated with a decline in naïve T cell numbers and a reduction in TCR diversity, which collectively reduce the immune system’s capacity to respond to novel antigens.^79^ Indeed, immunosenescence due to aging impairs the priming of T cells and the subsequent proliferation of primed T cells.^80-82^ The gradual decline in the production of naïve CD8^+^ T cells with age drives a shift in the population of memory CD8^+^ T cells toward exhaustion after repeated antigen exposure.^44^ This state of exhaustion renders T cells less responsive to new antigens and less effective in recognizing APCs.^83, 84^ In addition, the telomer shortening in CD8^+^ T cells can cause cell senescence by reducing proliferation and functionality.^85, 86^ Moreover, exhausted CD8^+^ T cells upregulated inhibitory receptors on the surface (i.e., PD-1), which can reduce the capacity of immune cells to recognize PD-L1-expressing cancer cells.^87-90^ We also demonstrated similar trends for the antitumoral efficacy of OVA-specific CD8^+^ T cells generated on-chip or off-chip, where the cytotoxicity of LNPoC-generated OVA-specific CD8^+^ T cells was significantly higher than off-chip. This is consistent with previous findings *in vivo*, demonstrating that the cytotoxic effect and effector functions of exhausted CD8^+^ T cells are limited and cannot effectively control the progression of tumors in the later stages.^87, 91, 92^

Our results revealed well-known trends in age-dependent decline in antigen presentation, T cell activation, and antitumoral efficacy characteristic of immunosenescence, which we further validated *in vivo* using the same species of mice. These age-dependent differences were apparent in our LNPoC but not in conventional 2D co-culture systems, suggesting that our LNPoC can uncover age-related immune function declines that are often masked in simplified culture systems. Unlike 2D models, which lack the spatial architecture and fluidic dynamics of the LN paracortex, the LNPoC facilitates continuous antigen flow and uptake, cell movement, and APC-T cell interactions within the culture microchamber due to the high surface-to-volume ratio inherent of OoC models.^93-96^ For example, T cell priming relies on dynamic spatial proximity and guided chemokine gradients to effectively interact, which are absent in static 2D cultures.^97^ In addition, biophysical cues can modulate immune cell functions, including macrophage polarization, T cell selection, and T cell activation.^98^ The LNPoC’s dynamic microenvironment provides physiologically relevant cell-cell interactions, where T cells can engage with APCs to mimic *in vivo* conditions. This design possibly improves the fidelity of antigen presentation and T cell activation by promoting a consistent flow and concentration gradient of antigens and signaling molecules, a feature that static cultures poorly replicate. Studies have shown that the microfluidic environment enables immune cells to respond to antigen stimuli with enhanced sensitivity and responsiveness by mimicking the natural APC-antigen and APC-T cell interactions that occur *in vivo*.^30^ Consequently, our LNPoC model is an accurate platform for studying immunosenescence, as it captures the nuanced cellular interactions and response dynamics that are critical to triggering age-related immune responses. Nevertheless, our LNPoC platform could be used to investigate immunosenescence effects on other vaccines, like influenza. This platform would require integrating additional immune cell types, such as CD4^+^ T cells, B cells, and plasma cells, to effectively capture germinal center dynamics and antibody production. However, accurately modeling CD8^+^ T cell priming and effector responses remains crucial in cancer immunotherapy, as these cells are key to tumor clearance after vaccination.

Despite the development of many OoC models recapitulating various functions of the LNs,^99^ none have focused on recapitulating immunosenescence on-a-chip. Most often, LN on-a-chip models are developed to mimic the architecture, biochemical environment, and fluid dynamics of actual LN and enable studies on immune cell behavior and interactions. The design of the microfluidic chip intends to capture different key functions of the LN and its complex role in immune responses.^30^ Several LN on-a-chip models recreate the distinct cellular regions of the LN, where different compartments allow the co-culture and spatial distribution of immune cells to study APC-T cell interactions and T cell activation.^100, 101^ These models can also replicate chemokine gradients to study immune cell chemotaxis, where immune cells migrate in response to controlled gradients of chemokines, similar to their *in vivo* counterparts.^102-104^ Several models have been used to test vaccines and to study the immunogenicity of antigens,^105, 106^ including some in an age-dependent manner.^107^ Indeed, immune cells from young and old human donors demonstrated differential age-associated responses to the influenza vaccine, where old immune cells exhibited reduced IgG production and CD4^+^ T cell activation upon vaccine stimulation compared to young immune cells.

Vaccine responses modeled on a chip primarily focus on the development of antibodies, with antigen presentation, T cell activation, germinal center formation, and antibody production.^108, 109^ In contrast, our LNPoC platform models CD8^+^ T cell-specific immune responses, which are the most important to evaluate following cancer vaccination against specific TAAs.^110, 111^ Also, our platform allows the collection of antigen-specific T cells for downstream application, particularly to assess the potency of effector functions downstream. Most importantly, our LNPoC was validated with the same species *in vivo*, which is a crucial step to ensure that the observed immune responses, particularly the age-dependent declines in antigen presentation, T cell activation, and antitumoral response, directly correlate with physiological outcomes. These *in vivo* validations are rarely carried out with the same species.^112^ This species discrepancy could hinder the translation of cancer vaccines to patients, with potential failures in clinical trials remaining.^113^ This same-species *in vivo* validation approach strengthens the translational relevance of our LNPoC model by confirming that it can accurately replicate the species-specific immune dynamics observed in live organisms. Moreover, it supports the LNPoC’s reliability in capturing age-related immune function changes and its potential as a predictive tool for evaluating immunosenescence and informing the design of clinical trials to test cancer vaccine efficacy.

Cancer vaccines, particularly those targeting CD8^+^ T cells, are promising immunotherapies for cancer treatment.^114^ However, their efficacy weakens in older populations due to immunosenescence.^115, 116^ In this context, our findings have several important implications for cancer vaccine development. First, the LNPoC is a valuable tool for the preclinical screening of cancer vaccines, allowing the testing of multiple candidates in an immune-relevant system before moving to costly and time-consuming clinical trials.^117^ By providing accurate age-dependent immune responses, our platform can potentially reduce the high failure rates of cancer vaccines in clinical trials and accelerate the development of more effective cancer immunotherapies. Specifically, the ability to model immunosenescence within our *in vitro* platform addresses a critical gap in the preclinical phase of current vaccine development pipelines.^118^ For aging populations, constituting the majority of cancer patients, there is an increasing need for effective vaccines;^43-45^ however, these populations are unaccounted for in preclinical models. Therefore, our LNPoC could provide insights into how vaccines can be optimized for older patients and lead to the development of new vaccine formulations or adjuvants specifically designed to enhance immune responses in geriatric populations. Lastly, our LNPoC platform could be a powerful tool for personalized medicine by enabling tailored testing of cancer vaccine efficacy for elderly patients from blood-isolated immune cells. Our LNPoC could guide the prediction of older individuals’ immune responses to various immunotherapies, which is invaluable for testing and optimizing a broader range of immunotherapies to develop more effective treatments for the aging population.

Despite its strengths, our LNPoC platform has several limitations. We used murine rather than human immune cells, which limits the direct applicability of findings to human systems due to interspecies differences in immune response characteristics.^119^ Nonetheless, our *in vivo* validation confirms that our platform can reliably recapitulate immune responses in an age-dependent manner. Furthermore, our LNPoC lacks extracellular matrix (ECM)-cell interactions, which are crucial for accurately mimicking cell migration and positioning, and integral to immune responses in lymphoid tissues.^120^ Additionally, the simplified tissue architecture of LNPoC, without stromal or follicular dendritic cells, may overlook critical cell-cell and cell-ECM interactions essential for a fully representative immune microenvironment.^121^ Our LNPoC is still a simplified model that cannot fully capture the complexity of the immune system. For instance, the platform focuses primarily on the lymph node paracortex and does not include other critical immune responses (i.e., memory or antibody), which also play important roles in vaccine-induced immune responses.^122^ Like other OoC technologies, our LNPoC represents an isolated environment, excluding systemic interactions with other immune organs, such as the spleen and bone marrow, which are vital for a holistic understanding of immune dynamics after vaccination.^123^ While the platform simulates some aspects of immunosenescence, additional research is needed to fully incorporate the range of systemic, cellular, and molecular changes that occur with aging.^74^ Although the current study used mice that were 35 to 36 weeks old to model early immunosenescence. Future research will focus on incorporating immune cells from mice aged 68 weeks or older. This will help further validate and expand the applicability of the LNPoC platform in modeling immune dysfunction associated with late-stage aging. We utilized the OVA-I peptide instead of full-length OVA protein for antigen presentation assays. While this peptide allows direct MHC-I-mediated antigen presentation and CD8^+^ T cell activation, it bypasses natural antigen processing. Future versions of the LNPoC platform will contain full-length proteins or patient-specific tumor antigens to assess better the complete pathways of antigen processing and presentation by APCs. Our *in vivo* validation demonstrated that young mice had significantly higher OVA-specific CD8+ T cell responses than older mice in blood, lymph nodes, and tumors. However, the differences in tumor size between the groups were not statistically significant over the 21-day observation period. This may be due to the aggressive nature of the B16-OVA tumor model, where rapid growth can outpace immune control efforts.^124, 125^ Additionally, immunosuppressive factors in the tumor microenvironment and the short study duration may have suppressed the significant differences in tumor burden, despite notable variations in immune activation.^125^ Future studies with more extended observation periods or less aggressive tumor models could illustrate the impact of immunosenescence on tumor control. Although we considered the tumor-killing ability of CD8^+^ T cells as “cytotoxicity,” our assessment was based on functional effector activity, measured by B16-OVA tumor cell viability assays after co-culture with OVA-specific CD8^+^ T cells. In this study, we did not directly measure cytotoxic molecules such as granzyme B or perforin. In future studies, we will incorporate granzyme B staining to more precisely characterize CD8^+^ T cell cytotoxic effector mechanisms and the corresponding pathways. In addition, future investigation of the LNPoC platform should incorporate clinically relevant neoantigens or influenza antigens to assess broader immunological responses beyond just CD8^+^ T cells. Another limitation of the current study is that the LNPoC platform utilized murine immune cells rather than human cells, which limits direct translational applicability. Due to significant physiological differences between murine and human immune systems, such as variations in antigen presentation and T cell receptor diversity, future studies will incorporate human PBMCs, dendritic cells, and T cells. The platform will also be optimized with human-specific cytokines, chemokines, and extracellular matrix components to simulate the human lymphoid microenvironment. Addressing these limitations by incorporating human cells, ECM components, and multi-organ system integration will be essential for furthering the utility and translational relevance of our LNPoC platform for cancer vaccine development and screening.

## Conclusions

Our LNPoC platform represents a significant advancement in studying the impact of immunosenescence on cancer vaccine discovery, development, and translation to the clinic. By replicating age-related declines in immune function, our LNPoC is a valuable tool for understanding the impact of immunosenescence on current cancer vaccines and guiding the development of more effective therapies for older populations. Our LNPoC platform is the first step in improving preclinical cancer vaccine testing by considering age as a variable. It could also be applied beyond cancer vaccines to other areas of immunotherapy, such as immune checkpoint inhibitors or adoptive cell therapies, where LN function plays a crucial role in activating immune cells. Continued research using this platform will be critical for achieving more personalized and age-specific cancer treatment strategies, ultimately improving therapeutic outcomes and quality of life for aging populations.

## Author contributions

Conceptualization: SM, AHN, VJ; Data curation: SM, AHN, SK, DK, CY, CJ, NM, VJ; Formal analysis: SM, AHN, VJ; Funding acquisition: AK, VJ; Investigation: SM, AHN, SK, DK, CY, CJ, NM, VJ; Methodology: SM, AHN, SK, DK, CY, CJ, NM, VJ; Supervision: MRD, AK, VJ; Visualization: SM, DK, VJ; Writing – original draft: SM, AHN, VJ; Writing – review & editing: SM, AHN, SK, DK, CY, CJ, NM, MRD, AK, VJ

## Acknowledgements

S.M. and A.H.N. contributed equally to this work. The authors acknowledge funding from the Terasaki Institute for Biomedical Innovation. We thank the NIH tetramer core for providing the OVA tetramer.

## References

1. J. Liu, M. Fu, M. Wang, D. Wan, Y. Wei and X. Wei, J Hematol Oncol, 2022, 15, 28.

2. M. Kaczmarek, J. Poznańska, F. Fechner, N. Michalska, S. Paszkowska, A. Napierała and A. Mackiewicz, Cells, 2023, 12.

3. M. Bilusic and R. A. Madan, Am J Ther, 2012, 19, e172–181.

4. Journal.

5. H. Donninger, C. Li, J. W. Eaton and K. Yaddanapudi, Vaccines, 2021, 9, 668.

6. A. Tojjari, A. Saeed, M. Singh, L. Cavalcante, I. H. Sahin and A. Saeed, Vaccines (Basel), 2023, 11.

7. H. Sajjad, S. Imtiaz, T. Noor, Y. H. Siddiqui, A. Sajjad and M. Zia, Animal Model Exp Med, 2021, 4, 87–103.

8. A. Loewa, J. J. Feng and S. Hedtrich, Nature Reviews Bioengineering, 2023, 1, 545–559.

9. A. D. Martin, X. Wang, M. L. Sandberg, K. R. Negri, M. L. Wu, D. Toledo Warshaviak, G. B. Gabrelow, M. E. McElvain, B. Lee, M. E. Daris, H. Xu and A. Kamb, Journal of Immunotherapy, 2021, 44, 95–105.

10. J. F. Vansteenkiste, B. C. Cho, T. Vanakesa, T. De Pas, M. Zielinski, M. S. Kim, J. Jassem, M. Yoshimura, J. Dahabreh, H. Nakayama, L. Havel, H. Kondo, T. Mitsudomi, K. Zarogoulidis, O. A. Gladkov, K. Udud, H. Tada, H. Hoffman, A. Bugge, P. Taylor, E. E. Gonzalez, M. L. Liao, J. He, J. L. Pujol, J. Louahed, M. Debois, V. Brichard, C. Debruyne, P. Therasse and N. Altorki, Lancet Oncol, 2016, 17, 822–835.

11. D. J. Schwartzentruber, D. H. Lawson, J. M. Richards, R. M. Conry, D. M. Miller, J. Treisman, F. Gailani, L. Riley, K. Conlon, B. Pockaj, K. L. Kendra, R. L. White, R. Gonzalez, T. M. Kuzel, B. Curti, P. D. Leming, E. D. Whitman, J. Balkissoon, D. S. Reintgen, H. Kaufman, F. M. Marincola, M. J. Merino, S. A. Rosenberg, P. Choyke, D. Vena and P. Hwu, New England Journal of Medicine, 2011, 364, 2119–2127.

12. G. A. Van Norman, JACC Basic Transl Sci, 2019, 4, 845–854.

13. D. Sun, W. Gao, H. Hu and S. Zhou, Acta Pharm Sin B, 2022, 12, 3049–3062.

14. O. Urzì, R. Gasparro, E. Costanzo, A. De Luca, G. Giavaresi, S. Fontana and R. Alessandro, Int J Mol Sci, 2023, 24.

15. C. Ma, Y. Peng, H. Li and W. Chen, Trends in Pharmacological Sciences, 2021, 42, 119–133.

16. J. J. Han, Artif Organs, 2023, 47, 449–450.

17. A. van Rijt, E. Stefanek and K. Valente, Cancers (Basel), 2023, 15.

18. Y. S. Zhang, J. Aleman, S. R. Shin, T. Kilic, D. Kim, S. A. Mousavi Shaegh, S. Massa, R. Riahi, S. Chae, N. Hu, H. Avci, W. Zhang, A. Silvestri, A. Sanati Nezhad, A. Manbohi, F. De Ferrari, A. Polini, G. Calzone, N. Shaikh, P. Alerasool, E. Budina, J. Kang, N. Bhise, J. Ribas, A. Pourmand, A. Skardal, T. Shupe, C. E. Bishop, M. R. Dokmeci, A. Atala and A. Khademhosseini, Proc Natl Acad Sci U S A, 2017, 114, E2293–e2302.

19. Q. Pi, S. Maharjan, X. Yan, X. Liu, B. Singh, A. M. van Genderen, F. Robledo-Padilla, R. Parra-Saldivar, N. Hu, W. Jia, C. Xu, J. Kang, S. Hassan, H. Cheng, X. Hou, A. Khademhosseini and Y. S. Zhang, Advanced materials (Deerfield Beach, Fla.), 2018, 30, e1706913–e1706913.

20. N. S. Bhise, J. Ribas, V. Manoharan, Y. S. Zhang, A. Polini, S. Massa, M. R. Dokmeci and A. Khademhosseini, Journal of controlled release : official journal of the Controlled Release Society, 2014, 190, 82–93.

21. S. R. Shin, T. Kilic, Y. S. Zhang, H. Avci, N. Hu, D. Kim, C. Branco, J. Aleman, S. Massa, A. Silvestri, J. Kang, A. Desalvo, M. A. Hussaini, S.-K. Chae, A. Polini, N. Bhise, M. A. Hussain, H. Lee, M. R. Dokmeci and A. Khademhosseini, Adv Sci (Weinh), 2017, 4, 1600522–1600522.

22. S. R. Shin, Y. S. Zhang, D.-J. Kim, A. Manbohi, H. Avci, A. Silvestri, J. Aleman, N. Hu, T. Kilic, W. Keung, M. Righi, P. Assawes, H. A. Alhadrami, R. A. Li, M. R. Dokmeci and A. Khademhosseini, Anal Chem, 2016, 88, 10019–10027.

23. M. Chernyavska, M. Masoudnia, T. Valerius and W. Verdurmen, Cancer Immunology, Immunotherapy, 2023, 1–13.

24. T. I. Maulana, E. Kromidas, L. Wallstabe, M. Cipriano, M. Alb, C. Zaupa, M. Hudecek, B. Fogal and P. Loskill, Advanced Drug Delivery Reviews, 2021, 173, 281–305.

25. A. L. Beckwith, L. F. Velásquez-García and J. T. Borenstein, Advanced Healthcare Materials, 2019, 8, 1900289.

26. S. Shim, M. C. Belanger, A. R. Harris, J. M. Munson and R. R. Pompano, Lab on a Chip, 2019, 19, 1013–1026.

27. X. Cui, C. Ma, V. Vasudevaraja, J. Serrano, J. Tong, Y. Peng, M. Delorenzo, G. Shen, J. Frenster, R.-T. T. Morales, W. Qian, A. Tsirigos, A. S. Chi, R. Jain, S. C. Kurz, E. P. Sulman, D. G. Placantonakis, M. Snuderl and W. Chen, eLife, 2020, 9, e52253.

28. J. M. Ayuso, S. Rehman, M. Virumbrales-Munoz, P. H. McMinn, P. Geiger, C. Fitzgerald, T. Heaster, M. C. Skala and D. J. Beebe, Science Advances, 2021, 7, eabc2331.

29. Y. Ando, E. L. Siegler, H. P. Ta, G. E. Cinay, H. Zhou, K. A. Gorrell, H. Au, B. M. Jarvis, P. Wang and K. Shen, Advanced healthcare materials, 2019, 8, 1900001.

30. A. Shanti, N. Hallfors, G. A. Petroianu, L. Planelles and C. Stefanini, Frontiers in Pharmacology, 2021, 12.

31. C. L. Slingluff, Jr., Cancer J, 2011, 17, 343–350.

32. M. A. J. Morsink, N. G. A. Willemen, J. Leijten, R. Bansal and S. R. Shin, Micromachines (Basel), 2020, 11.

33. A. E. R. Kartikasari, M. D. Prakash, M. Cox, K. Wilson, J. C. Boer, J. A. Cauchi and M. Plebanski, Front Immunol, 2018, 9, 3109.

34. L. Donoghue, K. T. Nguyen, C. Graham and P. Sethu, Micromachines (Basel), 2021, 12.

35. F. Groell, O. Jordan and G. Borchard, European Journal of Pharmaceutics and Biopharmaceutics, 2018, 130, 128–142.

36. S. N. Crooke, I. G. Ovsyannikova, G. A. Poland and R. B. Kennedy, Immunity & Ageing, 2019, 16, 25.

37. C. A. Chougnet, R. I. Thacker, H. M. Shehata, C. M. Hennies, M. A. Lehn, C. S. Lages and E. M. Janssen, The Journal of Immunology, 2015, 195, 2624–2632.

38. C. Wong and D. R. Goldstein, Curr Opin Immunol, 2013, 25, 535–541.

39. G. Del Giudice, J. J. Goronzy, B. Grubeck-Loebenstein, P.-H. Lambert, T. Mrkvan, J. J. Stoddard and T. M. Doherty, npj Aging and Mechanisms of Disease, 2017, 4, 1.

40. A. D. Foster, A. Sivarapatna and R. E. Gress, Aging health, 2011, 7, 707–718.

41. X. Li, C. Li, W. Zhang, Y. Wang, P. Qian and H. Huang, Signal Transduction and Targeted Therapy, 2023, 8, 239.

42. Y. Jiang, Y. Li and B. Zhu, Cell Death & Disease, 2015, 6, e1792–e1792.

43. B. Pereira, X.-N. Xu and A. N. Akbar, Frontiers in immunology, 2020, 11, 583019.

44. J. Lian, Y. Yue, W. Yu and Y. Zhang, Journal of Hematology & Oncology, 2020, 13, 151.

45. H. Donninger, C. Li, J. W. Eaton and K. Yaddanapudi, Vaccines (Basel), 2021, 9.

46. A. Agrawal and B. Weinberger, Frontiers in Aging, 2022, 3, 882494.

47. S. N. Crooke, I. G. Ovsyannikova, G. A. Poland and R. B. Kennedy, Immun Ageing, 2019, 16, 25.

48. W. Xu, G. Wong, Y. Y. Hwang and A. Larbi, Seminars in Immunopathology, 2020, 42, 559–572.

49. D. Weiskopf, B. Weinberger and B. Grubeck-Loebenstein, Transpl Int, 2009, 22, 1041–1050.

50. N. Mohaghegh, A. Ahari, C. Buttles, S. Davani, H. Hoang, Q. Huang, Y. Huang, B. Hosseinpour, R. Abbasgholizadeh, A. L. Cottingham, N. Farhadi, M. Akbari, H. Kang, A. Khademhosseini, V. Jucaud, R. M. Pearson and A. Hassani Najafabadi, ACS Nano, 2024, 18, 27764–27781.

51. N. Mohaghegh, A. Iyer, E. Wang, N. Z. Balajam, H. Kang, M. Akbari, M. S. Barnhill, A. Khademhosseini, R. M. Pearson and A. Hassani Najafabadi, Journal of Controlled Release, 2025, 382, 113670.

52. J. Kim, A. Morgott, Z. Wu, L. Hopaluk, M. Miles, W. Stoner and Q. Li, 2019.

53. M. Jafarkhani, Z. Salehi, M. A. Shokrgozar and S. Mashayekhan, International Journal for Numerical Methods in Biomedical Engineering, 2019, 35, e3154.

54. S. S. Jensen, H. Jensen, C. Cornett, E. H. Møller and J. Østergaard, Journal of Pharmaceutical and Biomedical Analysis, 2014, 92, 203–210.

55. S. Maity, T. Bhuyan, R. Bhattacharya and D. Bandyopadhyay, ACS Applied Materials & Interfaces, 2021, 13, 19430–19442.

56. S. Maity, T. Bhuyan, J. P. Pattanayak, S. S. Ghosh and D. Bandyopadhyay, Nanotechnology, 2021, 32, 505704.

57. F. Liao, J. Zhang, Y. Hu, A. H. Najafabadi, J. J. Moon, M. S. Wicha, B. Kaspo, J. Whitfield, A. E. Chang and Q. Li, Cancer Immunology, Immunotherapy, 2022, 71, 1959–1973.

58. A. H. Najafabadi, Z. I. N. Abadi, M. E. Aikins, K. E. Foulds, M. M. Donaldson, W. Yuan, E. B. Okeke, J. Nam, Y. Xu, P. Weerappuli, T. Hetrick, D. Adams, P. A. Lester, A. M. Salazar, D. H. Barouch, A. Schwendeman, R. A. Seder and J. J. Moon, Journal of Controlled Release, 2021, 337, 168–178.

59. J. Nikolich-Žugich, Nat Immunol, 2018, 19, 10–19.

60. Z. Zhou, Y. Pang, J. Ji, J. He, T. Liu, L. Ouyang, W. Zhang, X.-L. Zhang, Z.-G. Zhang, K. Zhang and W. Sun, Nature Reviews Immunology, 2024, 24, 18–32.

61. H. L. Thompson, M. J. Smithey, J. L. Uhrlaub, I. Jeftić, M. Jergović, S. E. White, N. Currier, A. M. Lang, A. Okoye, B. Park, L. J. Picker, C. D. Surh and J. Nikolich-Žugich, Aging Cell, 2019, 18, e12865.

62. C. Sebastián, C. Herrero, M. Serra, J. Lloberas, M. A. Blasco and A. Celada, J Immunol, 2009, 183, 2356–2364.

63. C. A. Chougnet, R. I. Thacker, H. M. Shehata, C. M. Hennies, M. A. Lehn, C. S. Lages and E. M. Janssen, J Immunol, 2015, 195, 2624–2632.

64. C. Sebastián, C. Herrero, M. Serra, J. Lloberas, M. A. Blasco and A. Celada, The Journal of Immunology, 2009, 183, 2356–2364.

65. S. Della Bella, L. Bierti, P. Presicce, R. Arienti, M. Valenti, M. Saresella, C. Vergani and M. L. Villa, Clin Immunol, 2007, 122, 220–228.

66. S. X. Leng, Q. L. Xue, J. Tian, Y. Huang, S. H. Yeh and L. P. Fried, Exp Gerontol, 2009, 44, 511–516.

67. C. E. Moss, H. Phipps, H. L. Wilson and E. Kiss-Toth, Frontiers in Immunology, 2023, 14.

68. E. Linehan and D. C. Fitzgerald, Eur J Microbiol Immunol (Bp), 2015, 5, 14–24.

69. W. B. Ershler, W. H. Sun, N. Binkley, S. Gravenstein, M. J. Volk, G. Kamoske, R. G. Klopp, E. B. Roecker, R. A. Daynes and R. Weindruch, Lymphokine Cytokine Res, 1993, 12, 225–230.

70. M. F. Neurath and S. Finotto, Cytokine Growth Factor Rev, 2011, 22, 83–89.

71. Y. Oishi and I. Manabe, npj Aging and Mechanisms of Disease, 2016, 2, 16018.

72. A. L. Thomas, P. C. Alarcon, S. Divanovic, C. A. Chougnet, D. A. Hildeman and M. E. Moreno-Fernandez, Front Aging, 2021, 2, 732414.

73. I. M. Rea, D. S. Gibson, V. McGilligan, S. E. McNerlan, H. D. Alexander and O. A. Ross, Front Immunol, 2018, 9, 586.

74. J. Guo, X. Huang, L. Dou, M. Yan, T. Shen, W. Tang and J. Li, Signal Transduction and Targeted Therapy, 2022, 7, 391.

75. A. Alberro, A. Iribarren-Lopez, M. Sáenz-Cuesta, A. Matheu, I. Vergara and D. Otaegui, Scientific Reports, 2021, 11, 4358.

76. N. Unver and F. McAllister, Cytokine Growth Factor Rev, 2018, 41, 10–17.

77. A. C. Shaw, in Handbook of Immunosenescence: Basic Understanding and Clinical Implications, eds. T. Fulop, C. Franceschi, K. Hirokawa and G. Pawelec, Springer International Publishing, Cham, 2019, DOI: 10.1007/978-3-319-99375-1_98, pp. 981–992.

78. Z. Liu, Q. Liang, Y. Ren, C. Guo, X. Ge, L. Wang, Q. Cheng, P. Luo, Y. Zhang and X. Han, Signal Transduction and Targeted Therapy, 2023, 8, 200.

79. J. J. Goronzy and C. M. Weyand, Nature Reviews Immunology, 2019, 19, 573–583.

80. E. Carrasco, M. M. Gómez de las Heras, E. Gabandé-Rodríguez, G. Desdín-Micó, J. F. Aranda and M. Mittelbrunn, Nature Reviews Immunology, 2022, 22, 97–111.

81. L. A. Callender, E. C. Carroll, E. A. Bober and S. M. Henson, Ageing Research Reviews, 2018, 47, 24–30.

82. I. J. Rodriguez, N. Lalinde Ruiz, M. Llano León, L. Martínez Enríquez, M. D. P. Montilla Velásquez, J. P. Ortiz Aguirre, O. M. Rodríguez Bohórquez, E. A. Velandia Vargas, E. D. Hernández and C. A. Parra López, Front Immunol, 2020, 11, 604591.

83. M. S. Abdel-Hakeem, S. Manne, J.-C. Beltra, E. Stelekati, Z. Chen, K. Nzingha, M.-A. Ali, J. L. Johnson, J. R. Giles, D. Mathew, A. R. Greenplate, G. Vahedi and E. J. Wherry, Nature Immunology, 2021, 22, 1008–1019.

84. P. Tonnerre, D. Wolski, S. Subudhi, J. Aljabban, R. C. Hoogeveen, M. Damasio, H. K. Drescher, L. M. Bartsch, D. C. Tully, D. R. Sen, D. J. Bean, J. Brown, A. Torres-Cornejo, M. Robidoux, D. Kvistad, N. Alatrakchi, A. Cui, D. Lieb, J. A. Cheney, J. Gustafson, L. L. Lewis-Ximenez, L. Massenet-Regad, T. Eisenhaure, J. Aneja, W. N. Haining, R. T. Chung, N. Hacohen, T. M. Allen, A. Y. Kim and G. M. Lauer, Nature Immunology, 2021, 22, 1030–1041.

85. M. Bellon and C. Nicot, Viruses, 2017, 9.

86. K. Shirakawa and M. Sano, Cells, 2021, 10, 2435.

87. W. Jiang, Y. He, W. He, G. Wu, X. Zhou, Q. Sheng, W. Zhong, Y. Lu, Y. Ding, Q. Lu, F. Ye and H. Hua, Front Immunol, 2020, 11, 622509.

88. C. Fenwick, V. Joo, P. Jacquier, A. Noto, R. Banga, M. Perreau and G. Pantaleo, Immunol Rev, 2019, 292, 149–163.

89. E. J. Wherry and M. Kurachi, Nat Rev Immunol, 2015, 15, 486–499.

90. J. L. Collier, S. A. Weiss, K. E. Pauken, D. R. Sen and A. H. Sharpe, Nature Immunology, 2021, 22, 809–819.

91. V. Decman, B. J. Laidlaw, T. A. Doering, J. Leng, H. C. Ertl, D. R. Goldstein and E. J. Wherry, J Immunol, 2012, 188, 1933–1941.

92. J. Reiser and A. Banerjee, J Immunol Res, 2016, 2016, 8941260.

93. P. Moura Rosa, N. Gopalakrishnan, H. Ibrahim, M. Haug and Ø. Halaas, Lab on a Chip, 2016, 16, 3728–3740.

94. M. Rothbauer, H. Zirath and P. Ertl, Lab on a Chip, 2018, 18, 249–270.

95. V. Ortseifen, M. Viefhues, L. Wobbe and A. Grünberger, Frontiers in Bioengineering and Biotechnology, 2020, 8.

96. A. Polini, L. L. del Mercato, A. Barra, Y. S. Zhang, F. Calabi and G. Gigli, Drug Discovery Today, 2019, 24, 517–525.

97. K. Poornima, A. P. Francis, M. Hoda, M. A. Eladl, S. Subramanian, V. P. Veeraraghavan, M. El-Sherbiny, S. M. Asseri, A. B. A. Hussamuldin, K. M. Surapaneni, U. Mony and R. Rajagopalan, Front Oncol, 2022, 12.

98. X. Zhang, T. H. Kim, T. J. Thauland, H. Li, F. S. Majedi, C. Ly, Z. Gu, M. J. Butte, A. C. Rowat and S. Li, Curr Opin Biotechnol, 2020, 66, 236–245.

99. Q. Wang, Y. Yang, Z. Chen, B. Li, Y. Niu and X. Li, Pharmaceutics, 2024, 16.

100. A. Shanti, B. Samara, A. Abdullah, N. Hallfors, D. Accoto, J. Sapudom, A. Alatoom, J. Teo, S. Danti and C. Stefanini, Pharmaceutics, 2020, 12.

101. N. Hallfors, A. Shanti, J. Sapudom, J. Teo, G. Petroianu, S. Lee, L. Planelles and C. Stefanini, Bioengineering (Basel), 2021, 8.

102. U. Haessler, Y. Kalinin, M. A. Swartz and M. Wu, Biomed Microdevices, 2009, 11, 827–835.

103. U. Haessler, M. Pisano, M. Wu and M. A. Swartz, Proc Natl Acad Sci U S A, 2011, 108, 5614–5619.

104. B. G. Ricart, B. John, D. Lee, C. A. Hunter and D. A. Hammer, J Immunol, 2011, 186, 53–61.

105. L. Radke, G. Sandig, A. Lubitz, U. Schließer, H. H. von Horsten, V. Blanchard, K. Keil, V. Sandig, C. Giese, M. Hummel, S. Hinderlich and M. Frohme, Bioengineering (Basel), 2017, 4.

106. T. Kraus, A. Lubitz, U. Schließer, C. Giese, J. Reuschel, R. Brecht, J. Engert and G. Winter, J Pharm Sci, 2019, 108, 2358–2366.

107. A. Dauner, P. Agrawal, J. Salvatico, T. Tapia, V. Dhir, S. F. Shaik, D. R. Drake, 3rd and A. M. Byers, Vaccine, 2017, 35, 5487–5494.

108. A. Purwada, S. B. Shah, W. Béguelin, A. August, A. M. Melnick and A. Singh, Biomaterials, 2019, 198, 27–36.

109. A. Purwada and A. Singh, Nat Protoc, 2017, 12, 168–182.

110. J. R. Veatch, N. Singhi, S. Srivastava, J. L. Szeto, B. Jesernig, S. M. Stull, M. Fitzgibbon, M. Sarvothama, S. Yechan-Gunja, S. E. James and S. R. Riddell, The Journal of Clinical Investigation, 2021, 131.

111. Y. Chen, D. Yu, H. Qian, Y. Shi and Z. Tao, Journal of Translational Medicine, 2024, 22, 394.

112. D. E. Ingber, Nature Reviews Genetics, 2022, 23, 467–491.

113. H. Maeng, M. Terabe and J. A. Berzofsky, Curr Opin Immunol, 2018, 51, 111–122.

114. J. Liu, M. Fu, M. Wang, D. Wan, Y. Wei and X. Wei, Journal of Hematology & Oncology, 2022, 15, 28.

115. C. E. DeSantis, K. D. Miller, W. Dale, S. G. Mohile, H. J. Cohen, C. R. Leach, A. Goding Sauer, A. Jemal and R. L. Siegel, CA Cancer J Clin, 2019, 69, 452–467.

116. J. Lian, Y. Yue, W. Yu and Y. Zhang, J Hematol Oncol, 2020, 13, 151.

117. C. Ma, Y. Peng, H. Li and W. Chen, Trends Pharmacol Sci, 2021, 42, 119–133.

118. T. Fan, M. Zhang, J. Yang, Z. Zhu, W. Cao and C. Dong, Signal Transduction and Targeted Therapy, 2023, 8, 450.

119. J. Mestas and C. C. Hughes, J Immunol, 2004, 172, 2731–2738.

120. C. Frantz, K. M. Stewart and V. M. Weaver, J Cell Sci, 2010, 123, 4195–4200.

121. J. G. Cyster and S. R. Schwab, Annu Rev Immunol, 2012, 30, 69–94.

122. F. Sallusto, A. Lanzavecchia, K. Araki and R. Ahmed, Immunity, 2010, 33, 451–463.

123. D. Huh, G. A. Hamilton and D. E. Ingber, Trends Cell Biol, 2011, 21, 745–754.

124. J. Oh, A. Magnuson, C. Benoist, M. J. Pittet and R. Weissleder, JCI Insight, 2018, 3.

125. C. Pettan-Brewer, J. Morton, R. Coil, H. Hopkins, S. Fatemie and W. Ladiges, Pathobiol Aging Age Relat Dis, 2012, 2.

