## Supporting Information for "Modeling Immunosenescence on-a-chip: a platform for cancer vaccine efficacy assessment"

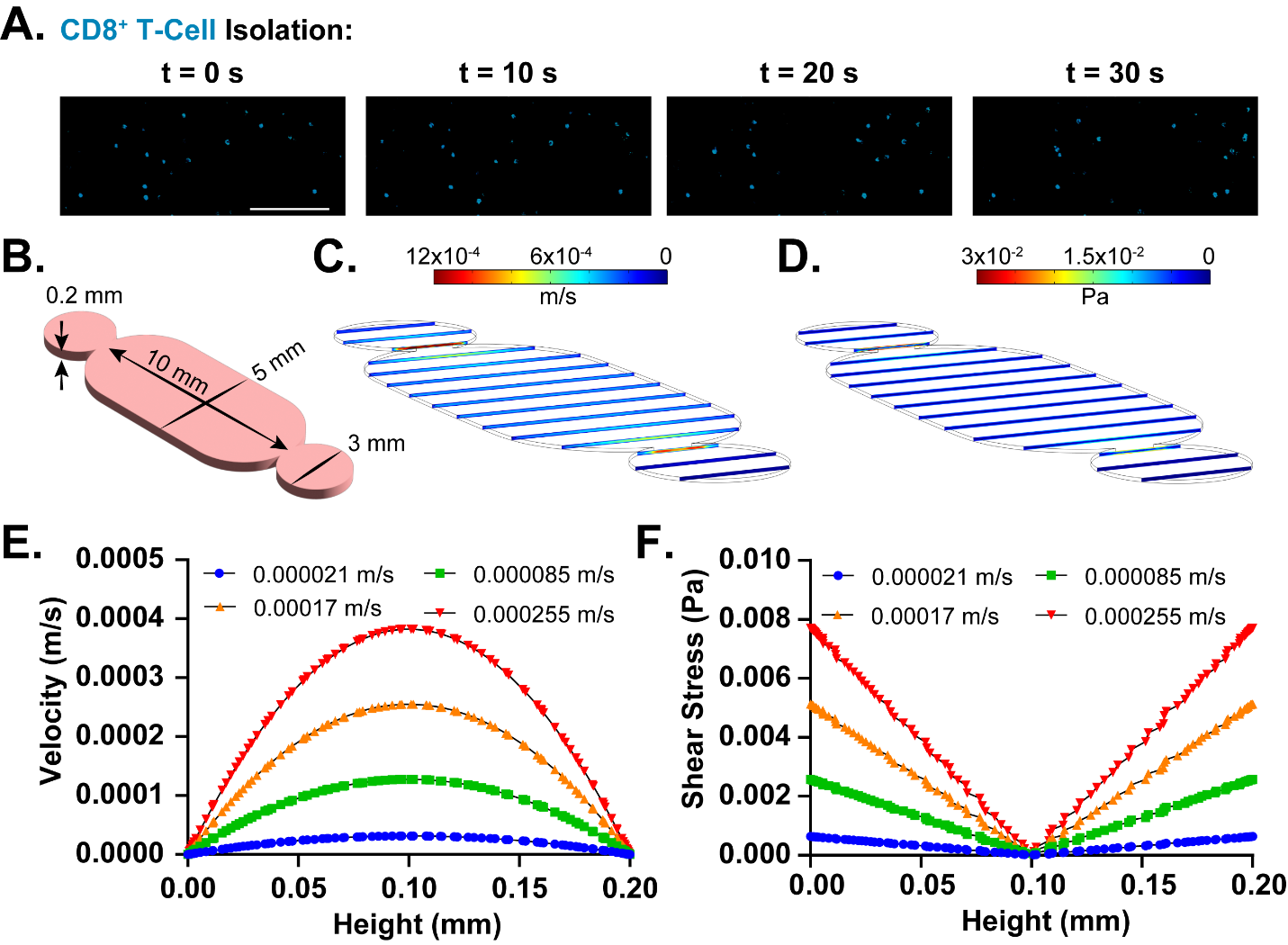

**Figure S1:** **CD8⁺ T-cell isolation and CFD analysis of flow behavior and shear stress within the LNPoC platform. A.** Fluorescence images showing isolated CD8⁺ T cells from the LNPoC device at an average velocity of 0.00017 m/s. Scale bar: 50 µm. **B.** Schematic representation of the LNPoC device geometry used for simulations. **C**-**D**. CFD simulation of velocity magnitude and shear stress distribution across the LNPoC device, showing flow distribution at an average velocity of 0.00017 m/s. **E**-**F**. Velocity profiles and shear stress distribution across the channel height for different inlet velocities (0.000021 m/s, 0.000085 m/s, 0.00017 m/s, and 0.000255 m/s).

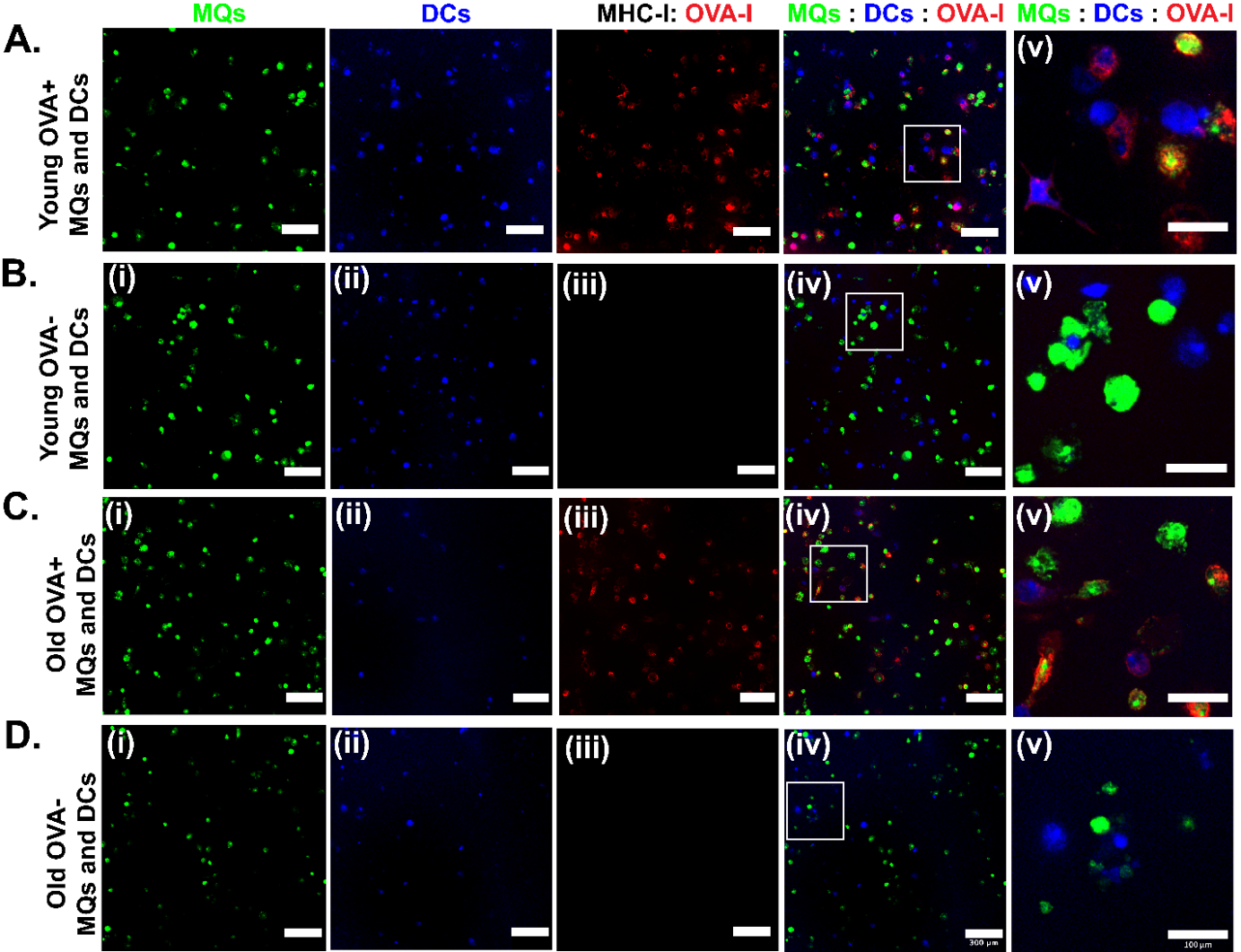

**Figure S2: Confocal analysis of on-chip presentation of OVA peptide by macrophages and dendritic cells. A.** Young MQs and DCs with OVA-I. **B.** Young MQs and DCs without OVA-I. **C.** Old MQs and DCs with OVA-I. **D.** Old MQs and DCs without OVA-I.

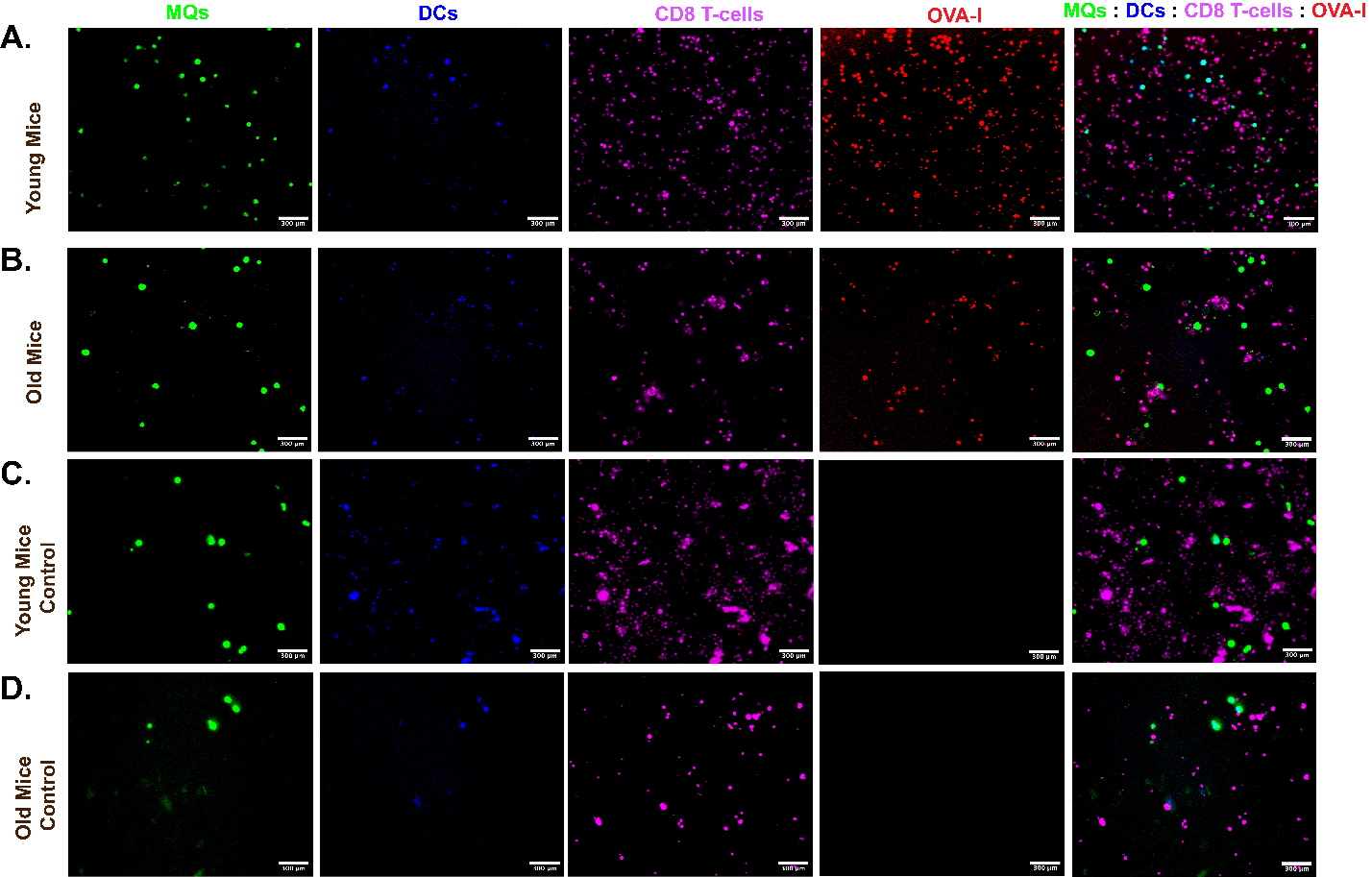

**Figure S3: Confocal analysis of on-chip activation of OVA-specific CD8^+^ T cells. A.** Young CD8^+^ T cell priming with OVA-I. **B.** Young CD8^+^ T cell priming without OVA-I. **C.** Old CD8^+^ T cell priming with OVA-I. **D.** Old CD8^+^ T cell priming without OVA-I.

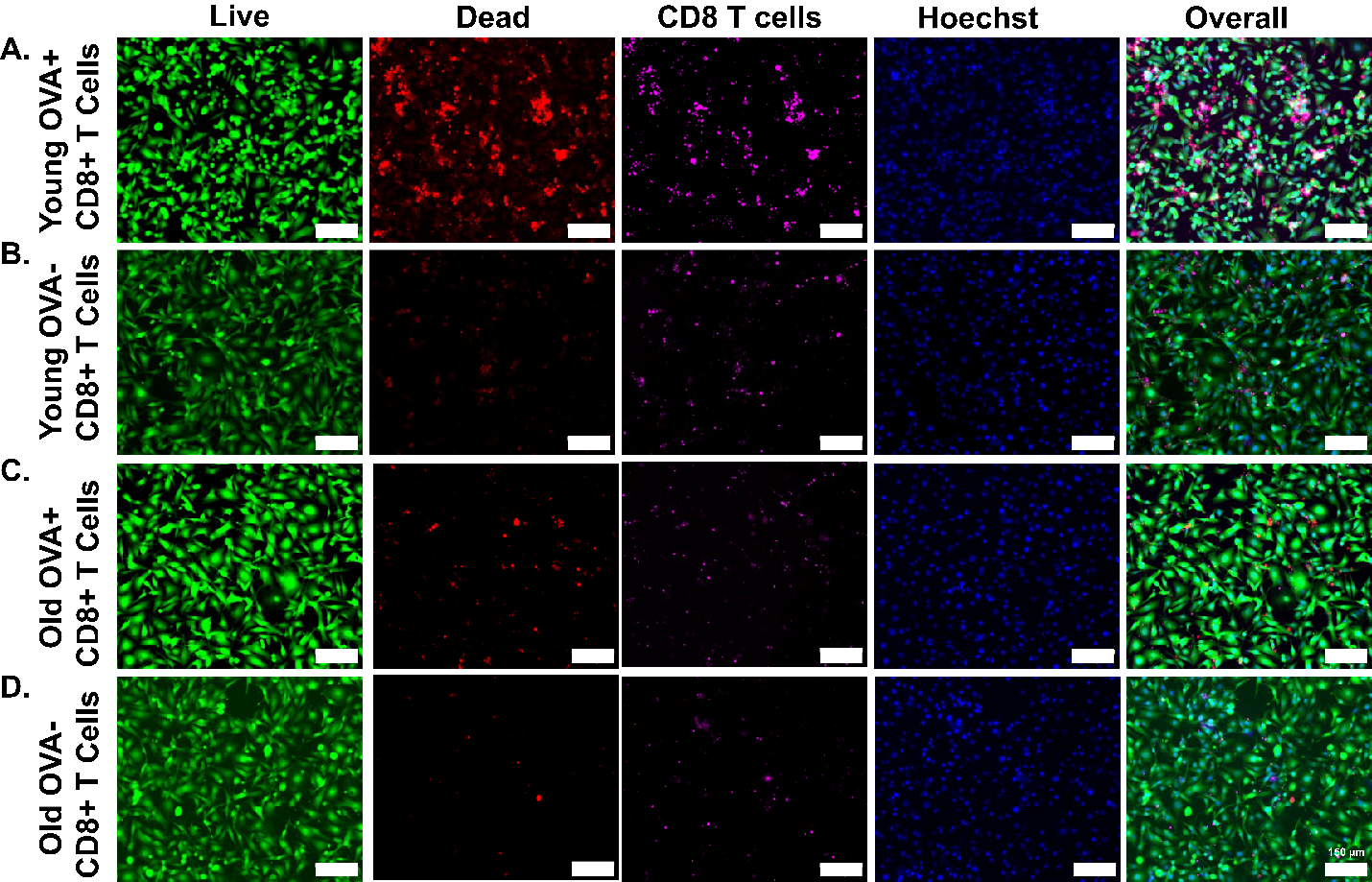

**Figure S4: Confocal analysis of antitumoral efficacy of OVA-I-specific CD8^+^ T cells.** **A.** Young CD8^+^ T cell priming with OVA-I. **B.** Young CD8^+^ T cell priming without OVA-I. **C.** Old CD8^+^ T cell priming with OVA-I. D. Old CD8^+^ T cell priming without OVA-I.

**Table S1: Comparison for OVA-specific CD8^+^ T cells:**

|  | **On-chip young CD8^+^ T cells** | **Young CD8^+^ T cells in the blood circulation at day 21.** | **Young CD8^+^ T cells in the LN at day 21.** |
| --- | --- | --- | --- |
| **Percentage of OVA-specific CD8^+^ T cells** | **12.3%** | **12.44%** | **13.35%** |
